# Suppressive Genetic Interactions Between Haploinsufficient Mitochondrial Genes Encoded in the 22q11.2 Microdeletion Locus Define Brain and Cardiac Phenotypes

**DOI:** 10.64898/2026.01.09.698677

**Authors:** Meghan Wynne, Stephanie A. Zlatic, Austin S. Park, Amanda Crocker, Hadassah Mendez-Vazquez, Eliana Liporace, Avanti Gokhale, Cristy Tower-Gilchrist, Maxine Robinette, Ryan H. Purcell, Gary J. Bassell, Erica Werner, Jennifer Q. Kwong, Victor Faundez

**Affiliations:** Department of Cell Biology, Emory University, Atlanta, GA, USA, 30322; Division of Pediatric Cardiology, Department of Pediatrics, Emory University School of Medicine, and Children’s Healthcare of Atlanta, Atlanta, GA, USA; Program in Neuroscience, Middlebury College, Bicentennial Way, Middlebury, VT 05753, United States; Nell Hodgson Woodruff School of Nursing. Emory University, Atlanta, GA, USA, 30322; Fralin Biomedical Research Institute at Virginia Tech Carilion, Roanoke, VA, USA 24016; School of Neuroscience, Virginia Tech, Blacksburg, VA, USA 24061

## Abstract

Genomic copy number variations, such as the 22q11.2 microdeletion syndrome, cause pleiotropic disorders that affect diverse organ systems and disrupt neurodevelopment. Deletions of the 22q11.2 locus reduce the dosage of up to 46 protein coding genes, raising questions about the identity of haploinsufficient genes and their genetic interactions contributing to 22q11.2 phenotypes. Here, we dissect functional and molecular relationships between two genes encoded within the 22q11.2 locus: the mitochondrial ribosomal protein gene MRPL40 and the mitochondrial citrate transporter SLC25A1. We show that a MRPL40 null mutation disrupts mitochondrial translation, impairs respiration, and affects multiple components of the SLC25A1 interactome including factors required for lipid metabolism, mitochondrial ribosome subunits, and the mitochondrial RNA processing machinery. *In silico* coessentiality network analysis revealed correlated and anticorrelated fitness interactions linking MRPL40 and SLC25A1 to mitochondrial translation, intermediate carbon metabolism, and interferon signaling. We determined that *Mrpl40*-null mutations are embryonic lethal in mice, but *Mrpl40^-/+^* mice are viable and displayed embryonic cardiac development and adult behavioral phenotypes. Similarly, *Slc25a1^+/-^* animals showed embryonic cardiac developmental defects but lacked the adult behavioral phenotypes observed in *Mrpl40^-/+^* mice. Surprisingly, transheterozygotic *Slc25a1^+/-^*;*Mrpl40^-/+^* mice suppressed or mitigated cardiac development, behavioral, and brain transcriptome phenotypes observed in single heterozygotic animals. These results reveal that MRPL40 and SLC25A1 are haploinsufficient genes within the 22q11.2 locus that genetically and biochemically interact to define tissue development and physiology. Our findings provide a framework for understanding the complexity and type of gene dosage interactions within the 22q11.2 deletion syndrome locus.

## Introduction

Human copy number variation (CNV) or microdeletion syndromes cause pleiotropic symptoms that affect multiple organ systems, including the brain, heart, immune and endocrine tissues, and craniofacial structures ^1–4^. This is exemplified by the 22q11.2 microdeletion syndrome, which can manifest as a constellation of multi-organ abnormalities or present primarily as a neurodevelopmental disorder, such as schizophrenia ^5–7^. The largest 22q11.2 microdeletion spans 3 Mb and reduces the dosage of 46 protein-coding genes, 27 pseudogenes, and 17 small non-coding regulatory RNAs ^8, 9^. Central questions arising from this genomic lesion include: which genes within the 22q11.2 CNV are dosage-sensitive contributors to disease phenotypes, and whether genes within the CNV genetically interact to specify these phenotypes? Current models consider the possibility of single dosage-sensitive “driver” genes within a CNV causing specific phenotypes; multiple dosage-sensitive genes contributing individually or through their genetic interactions; modifying loci residing outside the CNV influencing phenotypes; and imprinting mechanisms modulating gene effects ^10–12^. Genetic evidence in humans strongly supports the model that interactions among genes within and outside a CNV determine phenotypes ^10^. Indeed, *Drosophila* studies modeling genetic interactions among genes contained within either the 3q29 or 16p11.2 CNVs show that intra-CNV interactions can either enhance or suppress phenotypes ^13, 14^. To our knowledge, intra-CNV genetic interactions have not been systematically tested using engineered mutants in mice.

Modifier genomic loci outside the 22q11.2 CNV are known to influence phenotypes associated with this syndrome. These include cardiovascular defects that could be modulated by genes outside the 22q11.2 locus, such as either SLC2A3 or epigenetic regulator genes ^9, 15^. Likewise, psychiatric phenotypes may be modified by genes outside the 22q11.2 locus, whose function is required for mitochondrial homeostasis, including PARK2 and SPG7 ^16^. Notably, the 22q11.2 microdeletion is unique among CNVs because it removes the largest known group of nuclear-encoded genes required for mitochondrial function. These eight genes include *SLC25A1, TXNRD2, RTL10, MRPL40, PRODH, COMT, SNAP29*, and *TANGO2,* all localized to mitochondria according to Mitocarta 3.0 or new evidence ^17–20^. In addition, genes not classically annotated as mitochondrial may nevertheless influence mitochondrial biology, including *ZDHHC8, UFD1L,* and *DGCR8* ^17–19^.

The abundance of mitochondrial genes within the 22q11.2 locus and the prominent mitochondrial brain proteome phenotypes in 22q11.2 mouse models ^21, 22^ prompted us to test genetic interactions among 22q11.2 encoded mitochondrial genes. We asked if these mitochondrial genes could behave as haploinsufficient alleles and genetically interact to specify phenotypes in tissues with uniquely high mitochondrial bioenergetic demands, the brain and the heart ^23, 24^. We focused on two 22q11.2 mitochondrial genes *Slc25a1* and *Mrpl40*. *Slc25a1* encodes a citrate transporter located in the inner mitochondrial membrane that transports citrate from the mitochondrial matrix to the cytoplasm ^25, 26^. *Mrpl40* specifies a polypeptide belonging to the large 39S subunit of the mitochondrial ribosome that binds a structural Val-tRNA ^27, 28^. Slc25a1 and Mrpl40 proteins participate in a shared protein–protein interaction network ^29^. We have shown that like *Slc25a1*, *Mrpl40* is required for electron transport chain function and mitochondrial ribosome activity ^29, 30^. Because *Slc25a1* and *Mrpl40* converge on a mitochondrial protein interaction network, we predicted that their combined haploinsufficiency would exacerbate cardiac and brain phenotypes observed in the single-gene deficiencies. In contrast, and unexpectedly, we found that combined *Slc25a1* and *Mrpl40* haploinsufficiency suppressed cardiac and behavioral phenotypes observed in single haploinsufficiencies. These findings highlight the complexity of genetic interactions shaping CNV phenotypes and demonstrate the centrality of mitochondrial metabolism to heart development and behavior. We propose that, in addition to SLC25A1 and MRPL40, other dosage-sensitive nuclear-encoded mitochondrial genes contribute to the emergence of cardiovascular and psychiatric disease.

## Results

We confirmed our previously reported association between SLC25A1 and its interactome with MRPL40 ^29^, but this time using MRPL40 as a bait. We immunoprecipitated MRPL40 from detergent-soluble extracts of MRPL40-FLAG-expressing human neuroblastoma cells. MRPL40-FLAG coprecipitated endogenous SLC25A1 and proteins of the SLC25A1 interactome (Fig. 1A and Supplementary Fig. 1A). The specificity of these associations was tested by out-competition of immunoprecipitated antigens with an excess of FLAG peptide (Supplementary Fig. 1A, compare lanes 2-3). We then generated null alleles of MRPL40/*Mprl40* in human neuroblastoma cells and in the mouse germline to assess the robustness of biochemical and genetic interactions between SLC25A1 and MRPL40 ^29^. We resorted to engineer MRPL40 null cell lines because mouse mitochondrial ribosome subunit mutants of are lethal before gastrulation ^31^. SLC25A1-null mutations modify protein levels of mitochondrial ribosome subunits, RNA processing machinery, and lipid metabolism pathways belonging to the SLC25A1 interactome ^29^. Thus, we reasoned that a MRPL40 knock out should alter the levels of these SLC25A1 interactome proteins (Fig. 1A). We generated six MRPL40 null neuroblastoma clonal lines by CRISPR gene editing. We targeted exon one, which encodes the first 17 residues of MRPL40 (Supplementary Fig. 1A). We used these cell lines and their controls to quantify protein levels of SLC25A1 and its interactome (Fig. 1A). Loss of MRPL40 was confirmed by MRPL40 immunoblotting (Fig. 1B). MRPL40-null cells also showed severely reduced levels of mitochondrial ribosome translated MT-CO1, 2, 3, and MT-ATP6 (Fig. 1C, G and Supplementary Fig. 1B), and reduced abundance of mitochondrial ribosome subunits belonging to the SLC25A1 interactome (Fig. 1D,G). The decreased levels of these proteins resulted in a marked reduction of mitochondrial respiration, even after challenging cells with carbonyl cyanide 4-(trifluoromethoxy)phenylhydrazone or FCCP, as determined by Seahorse oximetry (Fig. 1H). MRPL40-null cells had an increased rate of media acidification, suggestive of increased glycolysis under severely compromised respiration (Fig. 1H). SLC25A1 protein levels did not change across MRPL40-null clones (Fig. 1B, G). However, proteins belonging to the SLC25A1 interactome involved in mitochondrial lipid metabolism (CPT1A) were significantly increased to 300%, across multiple MRPL40-null clones (Fig. 1E, G). Similarly, we observed modifications of SLC25A1 interactome proteins belonging to the mitochondrial RNA machinery in MRPL40-null cells (Fig. 1F-G). We found decreased levels of FASTKD2, TFAM, and PNPT1 to 60, 46, and 35%, respectively. These RNA processing machinery perturbations resulted in a 2-3-fold increase in the levels of the mitochondrial rRNAs RNR1 and RNR2 (Fig. 1I), 9 of the 13 mRNAs encoded in the mitochondrial DNA, and unprocessed RNA junctions, including those encoded in the light DNA strand (Fig. 1J). These increased RNA species resulted in an increased mitochondrial content of dsRNAs, an innate immunity inductor, detected with J2 antibody (Fig. 1K). These mitochondrial transcriptome alterations correlate with a doubling of the number of mitochondrial genomes in MRPL40-null cells (Fig. 1L). We conclude that MRPL40-null mutations alter the levels of components of the SLC25A1 interactome concomitant with broad alterations in the mitochondrial transcriptome. These findings reveal a wide spectrum of gain- and loss-of-function biochemical phenotypes in MRPL40-null mutations that extend beyond the function of the mitochondrial ribosome.

**Fig. 1.**
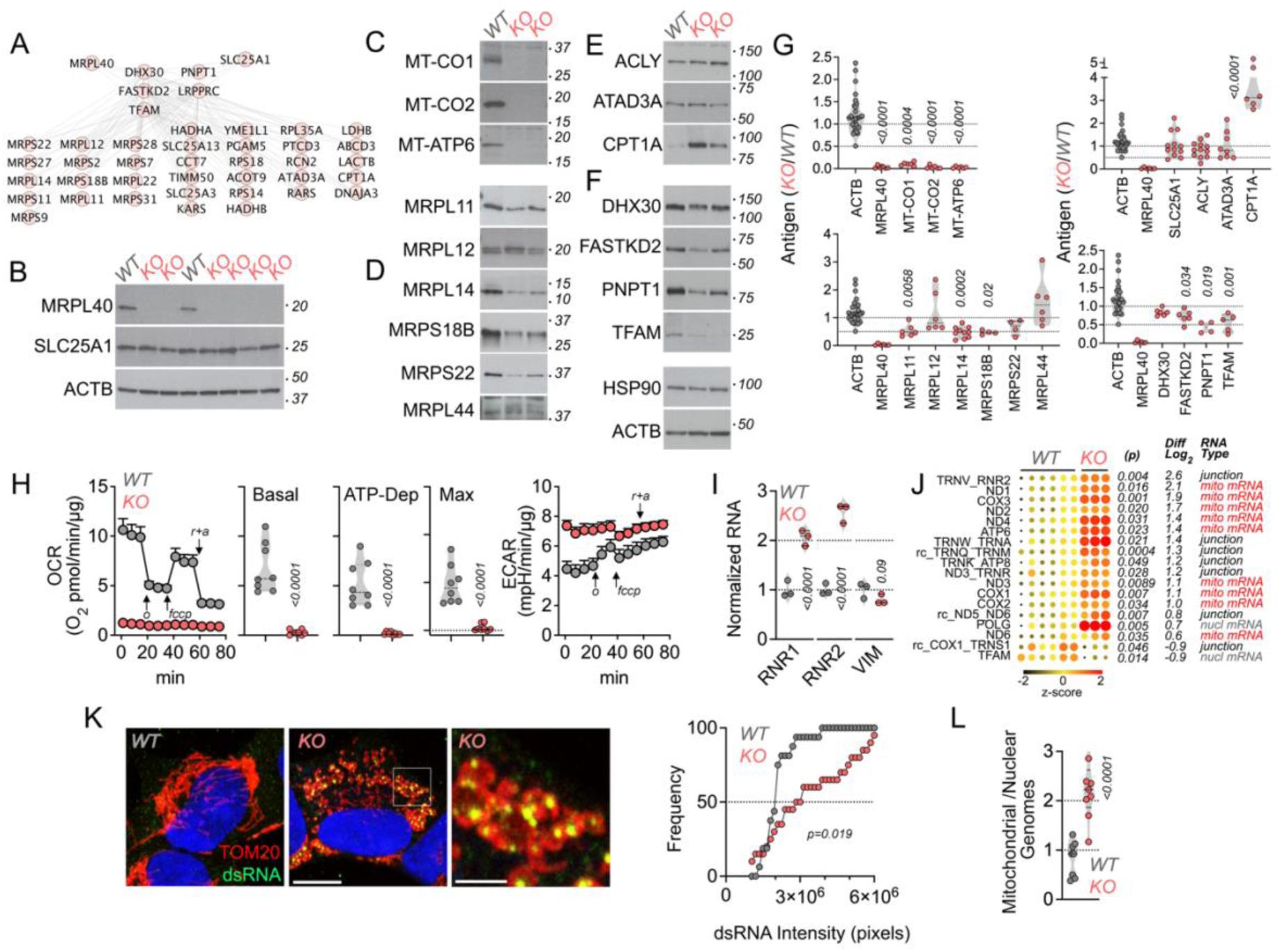
MRPL40 Null Mutants Disrupt Mitochondrial Ribosomes, Respiratory Chain, and Components of the SLC25A1 Interactome. **A.** Proteins of the SLC25A1 interactome according to Gokhale et al ^29^. Edges were defined by the proximity ligation mitochondrial interactome according to Antonicka et al. ^82^. **B.** MRPL40 and SLC25A1 immunoblots of wild type and MRPL40 null cell lines. **C.** Immunoblot of respiratory chain proteins encoded by the mitochondrial genome. **D.** Immunoblot of mitochondrial ribosome subunits. All subunits, except for MRPL44, are depicted in the SLC25A1 interactome. **E.** Immunoblot of SLC25A1 interactome proteins involved in acetyl-CoA and lipid metabolism. **F.** Immunoblot of RNA binding proteins present in the SLC25A1 interactome. HSP90 and ACTB were used as controls for B-F. **G**. Quantification of blots in panels B-F. Depicted is the ratio between mutant and wild type. Each dot represents an independent clone and/or independent experiment. p values were obtained with unpaired mean difference two-sided permutation t-test (italicized numbers represent p values). **H**. Metabolic activity in wild type and MRPL40 KO cells measured by the Seahorse Mito Stress Test (n=8 of each genotype). Oxygen consumption and extracellular acidification rates, OCR and ECAR, data are presented normalized to protein and analyzed by unpaired mean difference two-sided permutation t-test (italicized numbers represent p values). Basal, ATP-dependent, and maximal respiration were determined as described. Arrows indicate the sequential addition of oligomycin, FCCP and rotenone-antimycin. **I.** qRT-PCR quantification of mitochondrial rRNAs RNR1-2 and vimentin as control. Unpaired mean difference two-sided permutation t-test, n=3. **J**. Nanostring quantification of nuclear encoded and mitochondrial encoded RNAs. Junctions represent non-processed intermediaries derived from the polycistronic mitochondrial RNA. p values two-sided t test. Wild type n=6 and KO n=3. **K**. Immunofluorescent microscopy of with type and MRPL40 KO cells labeled for TOM20 and dsRNA. Scale bars correspond to 10 and 2.5 µm. Probability plot depicts the levels of dsRNA signal in 16 wild type and 20 MRPL40 KO cells. Kolmogorov-Smirnov test. **L**. Quantification of the ratio of mitochondrial and nuclear genomes. Unpaired mean difference two-sided permutation t-test. Wild type n=16 and KO n=20 cells.

The up- and down-regulation of components of the SLC25A1 interactome observed in MRPL40 KO cells suggests synthetic genetic interactions between the MRPL40 and SLC25A1 genes. Such interactions could either enhance and/or ameliorate phenotypes caused by deficiencies in either MRPL40 and/or SLC25A1 genes. We tested this prediction *in silico* by performing a coessentiality network analysis assessing genome-scale fitness correlations of SLC25A1 and MRPL40 across the publicly available Dependency Map (DepMap) screen datasets (Fig. 2) ^32, 33^. Correlated and anticorrelated genes reveal pathways and regulatory mechanisms connecting genes of interest. We performed this analysis using the FIREWORKS tool (Fitness Interaction Rank-Extrapolated netWORKs) ^34^. We identified two modules centered around MRPL40 and SLC25A1 that were connected by correlated and anticorrelated interactions with SOD2 and the zinc finger proteins ZNF577 and ZNF416 (Fig. 2A). These *in silico*-generated modules identified components of the SLC25A1 interactome that we determined experimentally ^29^, which were also altered in MRPL40 KO cells (red font, Fig. 2B). Within the MRPL40 module, the strongest positive fitness correlation was observed between MRPL40 and subunits of the mitochondrial ribosome and mitochondrial aminoacyl tRNA synthases (Fig. 2A, C). Similarly, the strongest positive fitness correlation within the SLC25A1 module included genes annotated to pyruvate and acetyl-CoA synthesis, which reside immediately upstream of citrate synthesis in the Krebs cycle. Importantly, the SLC25A1 and MRPL40 modules show anticorrelated genes involved in cytoplasmic protein synthesis (GO:0042273, RPL5;RPF2), nuclear encoded RNA processing (GO:0006396, LSM5;USP39), glycolipid biosynthesis (GO:0009247, PGAP2;ST8SIA1), and interferon signaling (GO:0060333, IRF1) (Fig. 2A, C). These *in silico* findings support the idea that these pathways could mediate SLC25A1 and MRPL40 genetic interactions in cells beyond the mitochondrial ribosome and citrate metabolism. Thus genetic interactions of similar complexity may occur in whole organisms where MRPL40 and SLC25A1 are simultaneously affected by a genetic defect.

**Fig. 2.**
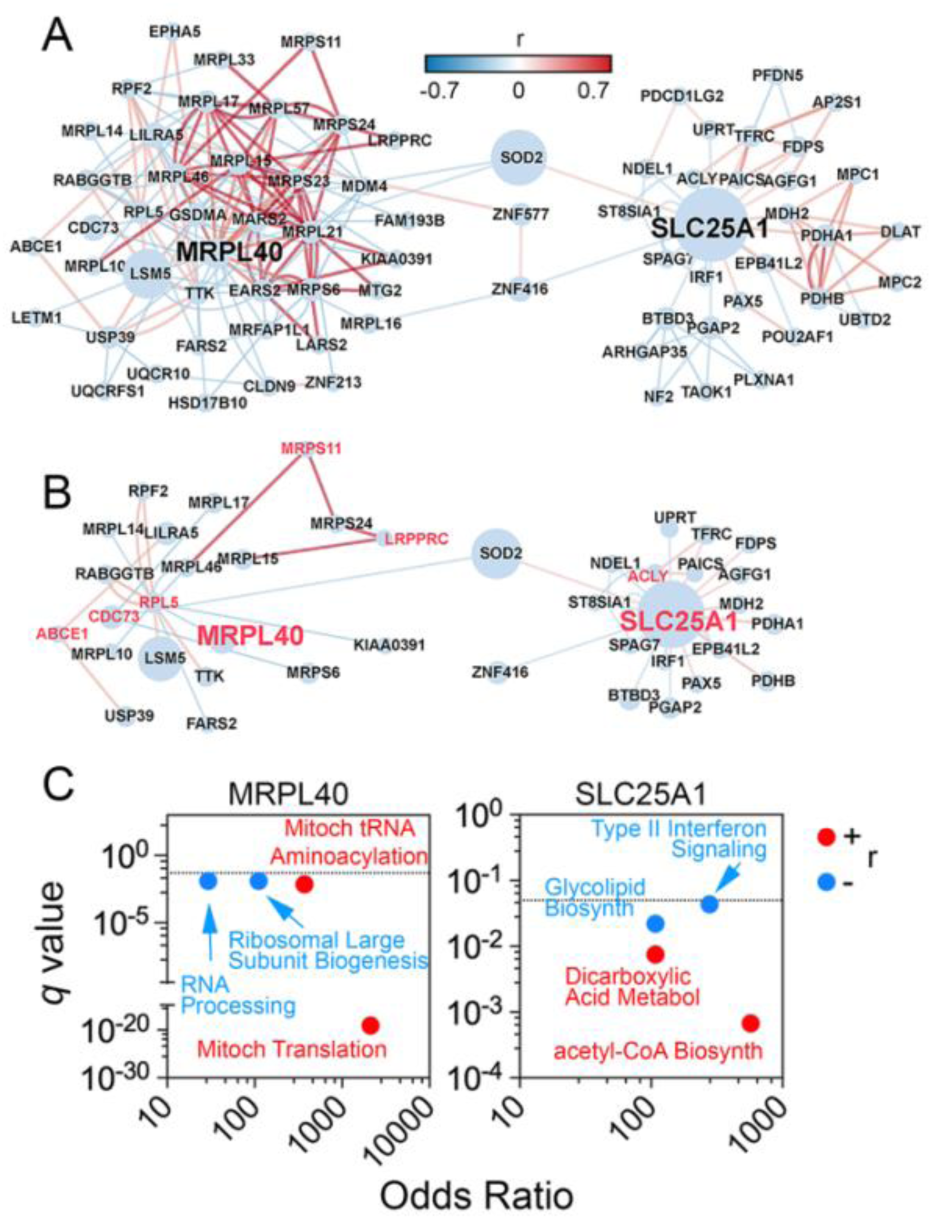
Coessentiality Network Analysis of SLC25A1 and MRPL40. **A.** Coessentiality network analysis showing genome-scale fitness correlations of SLC25A1 and MRPL40 as inputs. Nodes correspond to first and second level interaction nodes. Network of correlated (red edges) and anticorrelated genes (blue edges) was built with the FIREWORKS tool. r= Pearson correlation. **B**. Subnetwork of correlated and anticorrelated genes linked to components of the SLC25A1 interactome (red font nodes). **C**. ENRICHR Gene ontologies inferred from first order nodes connected to either SLC25A1 or MRPL40. Red and blue fonts depict correlated and anticorrelated genes, respectively.

The magnitude or quality of mouse phenotypes observed in single mutants of either the *Slc25a1* or *Mrpl40* genes could be modified in double mutant mice, a prediction founded on their protein interactions, mitochondrial functional convergence, and *in silico* predicted genetic interactions (Figs. 1 and 2). To generate a *Mrpl40*-null allele in mice, we first quantitatively estimated the essentiality of the mammalian MRPL40/*Mrpl40* gene by comparing the DepMap gene dependency score of human MRPL40 to TFAM and POLG, two proteins associated with the mitochondrial nucleoid ^32, 33, 35^ (Fig. 3A). We chose TFAM and POLG as their null alleles are lethal at or before mouse embryonic day 10.5 ^36, 37^. We used the pigmentation gene TYR/*Tyr*, as a reference since this gene is neither essential nor lethal in mammals ^38^. MRPL40 scored in between TFAM and POLG in essentiality, supporting lethality of homozygous *Mrpl40* null mutations before birth in mice (Fig. 3A). This prediction agrees with the pre-gastrulation developmental stalling observed in embryos carrying null mutations of other mitochondrial ribosome subunit genes ^31^. We generated a mouse *Mrpl40* mutant allele by CRISPR gene editing. We engineered the *Mrpl40* mutation in the heterozygotic *Slc25a1^-/+^*mouse strain we previously described to avoid genetic background variables when testing *Slc25a1* and *Mrpl40* transheterozygotic genetic interactions ^30^ (Fig. 3B). *Mrpl40* editing resulted in a 7bp deletion in exon 3, which is shared by the two annotated *Mrpl40* transcripts ENSMUST00000023391.16 (UNIPROT Q3UKS6) and ENSMUST00000119273.2 (UNIPROT D3Z7C0) (Fig. 3C). The founder *Mrpl40* mutant mouse carries a premature stop codon after residue 53, thus truncating the remaining 153 residues, or 75%, of the Mrpl40 primary sequence encoded by transcript ENSMUST00000023391.16 (Fig. 3C). The mutant stop codon resides more than 50-55 nucleotides upstream of the final exon-exon junction making it unlikely to undergo nonsense-mediated mRNA decay ^39^ (see Fig. 6C and Supplementary Fig 3C). The deleted primary sequence contains a domain that binds a structural Val-tRNA present in the large subunits of the mitochondrial ribosome ^27^. *Mrpl40^-/+^* animals were backcrossed at least 6 times to remove any off-target CRISPR edits (Fig. 3B). We genotyped animals at embryonic date 14.5, recovering 50% of the animals either as wild type or *Mrpl40^-/+^* (Fig. 3D). We did not obtain *Mrpl40^-/-^* embryos, indicative of lethality before embryonic date 14.5, as predicted by the gene dependency score (Fig. 3D). In contrast, we obtained all four genotypes at the expected Mendelian ratios in intercrosses of *Slc25a1^-/+^* with *Mrpl40^-/+^* animals (Fig. 3E). We did not observe any gross anatomical alterations in transheterozygotic *Slc25a1^-/+^*;*Mrpl40^-/+^* embryos and the size of newborn *Slc25a1^-/+^*;*Mrpl40^-/+^*animals was normal as compared with the wild type and their heterozygotic littermates.

**Fig. 3.**
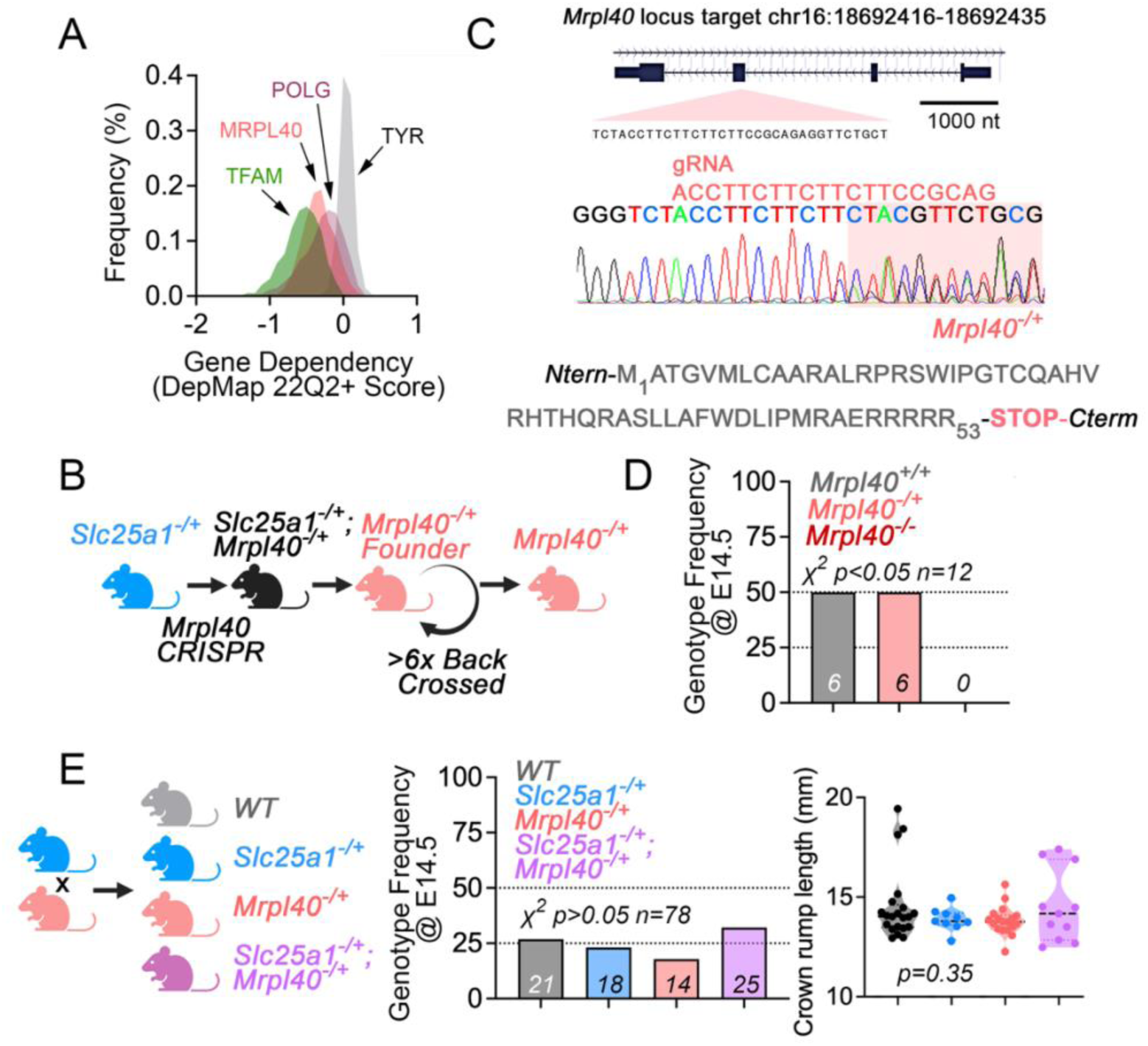
Generation of an *Mrpl40* mutant mouse. **A.** Estimation of mouse embryonic viability using DepMap gene dependency scores of human MRPL40, TFAM, POLG, and TYR. **B**. Experimental design to generate an *Mrpl40* null allele. **C**. *Mrpl40* genomic region, CRISPR mutagenesis strategy, founder mutation, and predicted protein truncation. **D**. Frequence of *Mrpl40* genotypes at embryonic age 14.5 from timed *Mrpl40^-/+^* intercrosses. **E**. Diagram of *Slc25a1^-/+^*;*Mrpl40^-/+^* timed intercrosses, frequency of genotypes, and crown-rump lengths of embryos at embryonic age 14.5. Chi square test of expected and observed genotypes.

We took advantage of the fact that our *Slc25a1^-/+^* mouse model shows haploinsufficient cardiac development and metabolic phenotypes to test whether *Slc25a1* and *Mrpl40* genetically interact to modulate tissue phenotypes ^30^. As previously described, *Slc25a1^-/+^*mice show ventricular septal defects and ventricular noncompaction revealed by decreased compact myocardium and increased trabecular myocardium ^30^(Fig. 4). Some of these phenotypes are common with the 22q11.2 microdeletion, which causes conotruncal and ventricular septal defects ^6^. We measured these *Slc25a1^-/+^* cardiac phenotypes in *Mrpl40^-/+^* and transheterozygotic *Slc25a1^-/+^*;*Mrpl40^-/+^* hearts. We chose to study hearts at embryonic day 14.5 when septation and ventricular wall compaction should be complete ^40, 41^. We recapitulated *Slc25a1^-/+^* septal defects in *Mrpl40^-/+^*at a similar frequency as compared to *Slc25a1^-/+^* animals (20-31.3%, X^2^=0.47, Fig. 4A-B, arrows). However, the frequency of *Slc25a1^-/+^* septal defects was significantly reduced to 5.6% in *Slc25a1^-/+^*;*Mrpl40^-/+^*mice (X^2^=0.049, Fig. 4A-B). This suppressive transheterozygotic interaction was also observed in the ventricular noncompaction phenotype (Fig. 4A and C, asterisks). We found a significant decrease in compact myocardium with a concomitant increase in the trabecular myocardium in *Slc25a1^-/+^* hearts but not in the *Mrpl40^-/+^* ventricular wall (Fig. 4A and C, asterisks). Importantly, the non-compaction phenotype observed in *Slc25a1^-/+^* hearts was ameliorated in transheterozygotic *Slc25a1^-/+^*;*Mrpl40^-/+^*mice (Fig. 4A and C, asterisks). These results show partially overlapping cardiac development phenotypes in single heterozygotic mice, which can be reverted closer to wild type in transheterozygotic *Slc25a1^-/+^* and *Mrpl40^-/+^* tissue.

**Fig. 4.**
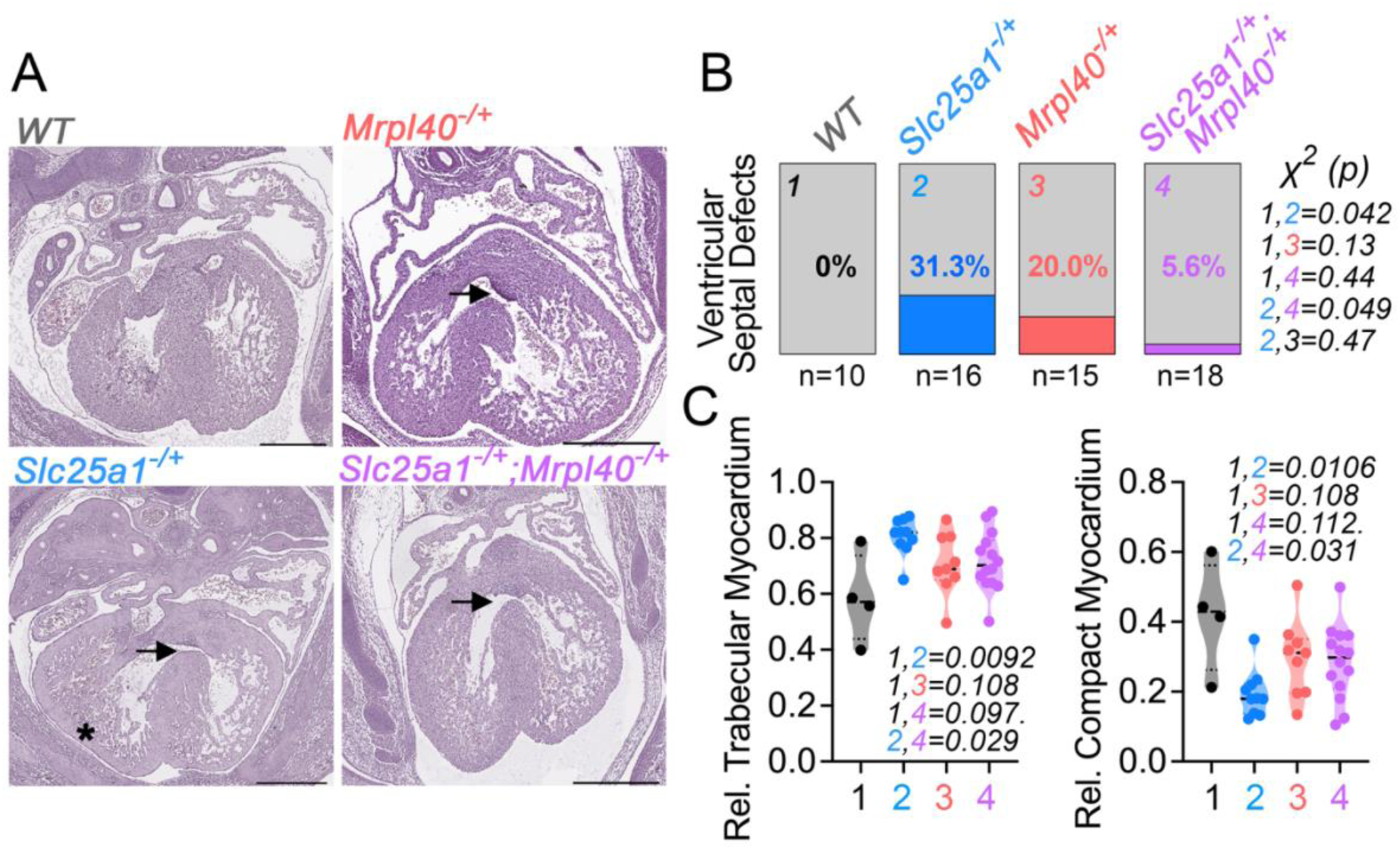
*Slc25a1^-/+^*;*Mrpl40^-/+^* Heterozygotic Mice Suppress Cardiac Defects Observed in Single Heterozygotes. **A**. Representative images of E14.5 embryos hearts from the indicated genotypes stained with hematoxylin and eosin. Asterisk indicates noncompaction and arrow indicates ventricular septal defect. Scale bar is 0.5 mm. **B** Frequency of cardiac defects observed in E14.5 in embryos with the indicated genotypes. Chi square test of expected and observed genotypes. **C** Quantification of relative compact myocardium thickness. one-way ANOVA Fisher’s LSD.

*Slc25a1^-/+^* and *Mrpl40^-/+^* transheterozygotic genetic interactions observed in hearts could expand to other organs affected in 22q11.2 microdeletion syndrome, such as the brain ^6^. Thus, we performed a battery of behavioral tests and collected microdissected adult brain cortex and hippocampus for bulk RNAseq from the same animals. We conducted Morris water maze (Fig. 5A), elevated plus maze, marble burying, nestlet shredding (Fig. 5B), light-dark cycle activity (Fig. 5C), and pre-pulse inhibition (Fig. 5D) to capture a wide span of behaviors in male and female mice (Supplementary Fig. 2). The Morris water maze, elevated plus maze, marble burying, and nestlet shredding were normal in all the genotypes tested irrespective of sex. Instead, we found that *Mrpl40^-/+^* males, but not females, have altered light-dark cycle activity with a bout of hyperactivity at dawn (mixed-effect model, genotype p=0.038, Fig. 5C). This phenotype was absent in *Slc25a1^-/+^* mutants as well as transheterozygotic *Slc25a1^-/+^*;*Mrpl40^-/+^* males supporting a suppressive genetic interaction between *Slc25a1^-/+^* and *Mrpl40^-/+^*. We further tested this finding by analyzing data with two-sided permutation statistics, which confirmed the phenotype in *Mrpl40^-/+^* mice whereas transheterozygotes were not different from wild type animals.

**Fig. 5.**
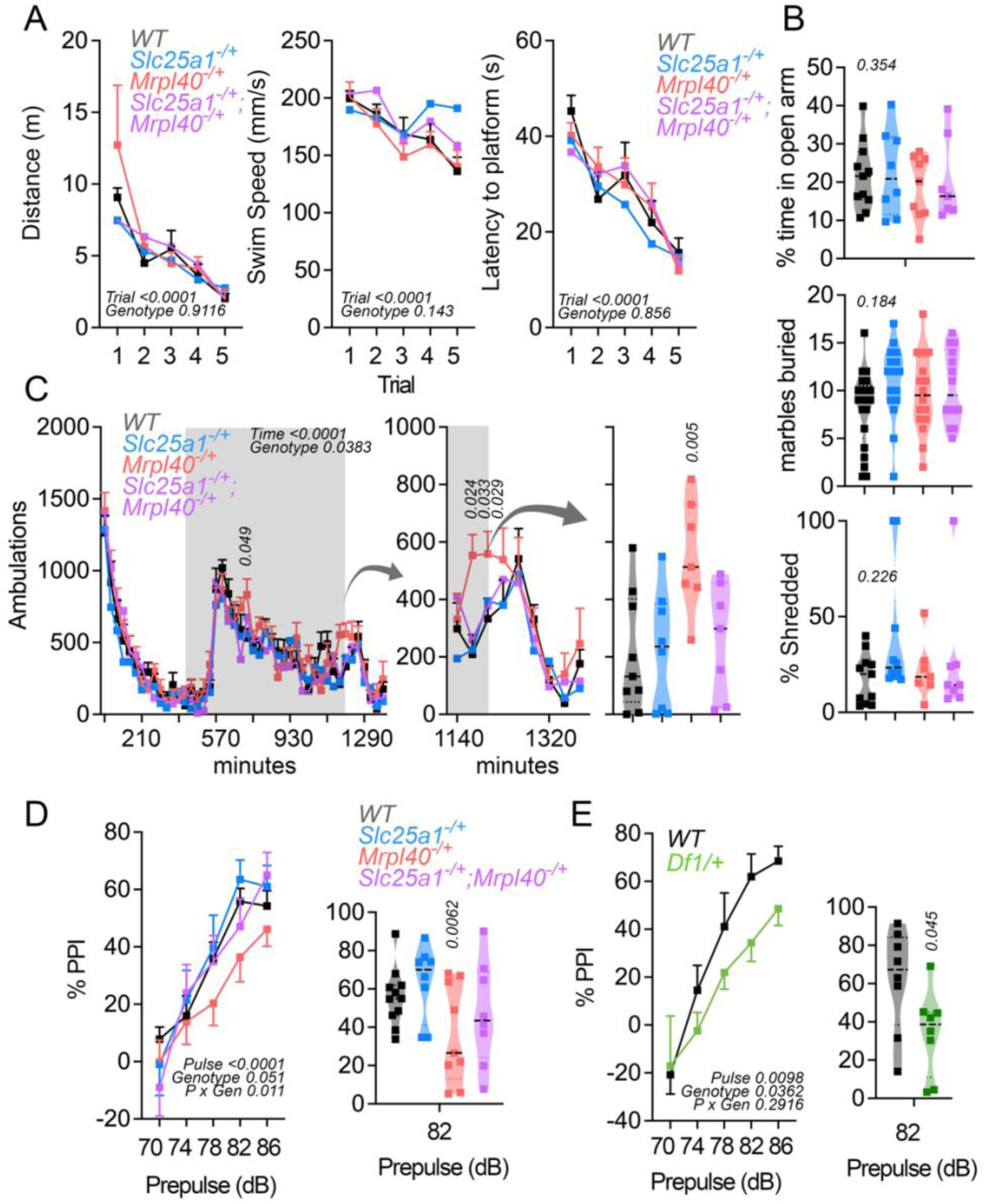
*Slc25a1^-/+^*;*Mrpl40^-/+^* Heterozygote Male Mice Suppress *Mrpl40^-/+^* Behavioral Phenotypes. **A**. Swim distance, swim speed, and latency to the platform during acquisition trials of the Morris water maze in 8-week-old males of the indicated genotypes. Two Way Repeated Measures ANOVA n=5 per genotype. **B.** Anxiety-like behaviors tests (elevated plus maze, marble burying, and nestlet shredding) are normal in all genotypes tested. One Way ANOVA followed by Holm-Šídák’s multiple comparisons test. n=7-11 per genotype. **C**. Time course of 23-h locomotor activity of animals of the specified genotype. Grey column denotes dark period. Data represent mean ± SEM. Statistical comparisons were performed using a mixed-effects model (Restricted Maximum Likelihood) with factors genotype, time, and their interactions. Mixed effect model was followed by either Fisher’s LSD or two-sided permutation t-tests at the indicated time points. Violin plot shows two-sided permutation t-tests of one of those points. n=7-9 per genotype. **D**. Percent PPI, data represent mean ± SEM. Statistical comparisons were performed using a mixed-effects model (Restricted Maximum Likelihood) with factors genotype, prepulse intensity, and their interactions. Mixed effect model was followed by two-sided permutation t-tests. Violin plot shows two-sided permutation t-tests at 82 dB. n=8-11 per genotype. **E**. Percent PPI as in D. Two Way Repeated Measures ANOVA n=8. Factors are genotype, prepulse intensity, and their interactions. Violin plot shows two-sided permutation t-test at 82 dB. n=8.

A well-established phenotype in mouse models of 22q11.2 microdeletion syndrome is an impaired pre-pulse inhibition response, which has construct validity for abnormal sensorimotor gating responses in 22q11.2 microdeletion human subjects ^5, 6, 42^. We compared the effects of single and double heterozygosity in *Slc25a1* and *Mrpl40* (Fig. 5D) and compared responses in these animals to those in the *Del(16Es2el-Ufd1l)217Bld* (heretofore referred as *Df1/+*, Fig. 5E) mouse, an animal model of the 22q11.2 microdeletion syndrome that includes a copy number variation in the *Slc25a1* gene ^43, 44^. We used these *Df1/+* mice as a positive control and a way to assess phenotypic similarities between *Df1/+* with either single or transheterozygotic animals. We found decreased pre-pulse inhibition in *Mrpl40^-/+^* mice as determined by the modification of the pulse response by the *Mrpl40^-/+^* genotype (mixed-effect model, pulse-genotype p=0.011, Fig. 5D). The effect of the *Mrpl40^-/+^* genotype was near significance (mixed-effect model, genotype p=0.051). Thus, we further scrutinized the pre-pulse inhibition phenotype in *Mrpl40^-/+^* animals using two-sided permutation statistics across pulse levels (Fig. 5D). This analysis showed strong and significant effects in *Mrpl40^-/+^* mice at 82 dB, which was reverted to wild type levels in transheterozygotic *Slc25a1^-/+^*;*Mrpl40^-/+^* males. The extent of pre-pulse inhibition in *Mrpl40^-/+^* mice was comparable to the phenotype in *Df1/+* mice (mixed-effect model, genotype p=0.036, Fig. 5E). Similar to the *Mrpl40^-/+^* mutants, the most robust *Df1/+* phenotype was observed at 82 dB after two-sided permutation statistics.

*Slc25a1^-/+^* and *Mrpl40^-/+^* transheterozygotic genetic interactions at the behavioral level should correlate with modifications of gene expression. We measured gene expression by bulk RNAseq in microdissected adult brain cortex and hippocampus of wild type, single, and transheterozygotic mice to test this hypothesis prediction (Fig. 6 and Supplementary Fig. 3). Non-parametric Uniform Manifold Approximation and Projection (UMAP) of the whole bulk transcriptome revealed no separation across all genotypes suggesting discrete gene expression changes across genotypes. However, thresholding bulk transcriptomes by p<0.01 (one-way ANOVA) showed genotype clustering where the cortical *Mrpl40^-/+^* transcriptome and the hippocampal *Slc25a1^-/+^* transcriptome segregated from the other genotypes (Fig. 6A). Parametric data dimensionality reduction and Euclidean distance clustering (PCA) showed wild type, *Slc25a1^-/+^*; and *Slc25a1^-/+^*;*Mrpl40^-/+^*cortical transcriptomes co-clustered away from the *Mrpl40^-/+^* transcriptome in cortex (Fig. 6B). Similarly, wild type and *Slc25a1^-/+^*;*Mrpl40^-/+^*hippocampal transcriptomes segregated from the *Slc25a1^-/+^* transcriptome (Supplementary Fig. 3B). These changes in the *Mrpl40^-/+^* transcriptome were evident even though we could not detect changes in the expression of Mrpl40 transcripts in both brain regions, likely the result of the discrete *Mrpl40* 7bp deletion sparing the mutant transcript from non-sense mediated decay (Fig. 6C). In contrast, *Slc25a1^-/+^* and *Slc25a1^-/+^*;*Mrpl40^-/+^*transheterozygotic cortex and hippocampus showed a mean reduction in the *Slc25a1* mRNA to ∼60% of wild type (Fig. 6C). We selected differentially expressed mRNA by a fold of change >1.5 and a p value <0.01 (Welsh t-test) to identify less than 100 RNAs affected in either single or transheterozygotes as compared to wild type tissue (Fig. 6D-E). Of these RNAs, less than 40% were protein coding RNAs while the rest were non-coding transcripts (Fig. 6D). The paucity of differentially expressed coding RNAs prevented us from identifying pathways and ontologies enriched in these transcripts with either GSEA or ENRICHR tools. Importantly, we found that differentially expressed mRNAs in either *Slc25a1^-/+^* or *Mrpl40^-/+^* were attenuated or reverted to wild type levels in transheterozygotic cortex (Fig. 6E), as exemplified by the transcripts encoding Klra2, Lrrc69, Mhrt, and Oas2 in cortex (Fig. 6F) and Asb17, Ifit1bl2, and Ppp1r1c in hippocampus (Supplementary Fig. 3F). In contrast, other mRNA phenotypes became overt in transheterozygotic cortex, as illustrated by Car13, Ccr2, and Hsd17b13 mRNAs (Fig. 6G). These results show a complex transcriptional response where we observed two responses; first, we find differentially expressed cortical transcripts in single heterozygotic mutants that can be reverted to wild type levels in transheterozygotic *Slc25a1^-/+^*;*Mrpl40^-/+^*cortex. A second response is an additive phenotype in transheterozygotic cortical tissue. We demonstrate that behavioral phenotypes and some differentially expressed transcripts observed in either *Mrpl40^-/+^* or *Slc25a1^-/+^* mutants can be suppressed in transheterozygotic *Slc25a1^-/+^*;*Mrpl40^-/+^*mice. We conclude that *Slc25a1^-/+^* and *Mrpl40^-/+^*are haploinsufficiencient mitochondrial genes that genetically interact with a complex pattern of phenotypic outputs, some of which are suppressive. Our findings suggest that these and other genes within copy number variation loci interact to modulate the quality and magnitude of tissue phenotypes.

**Fig. 6.**
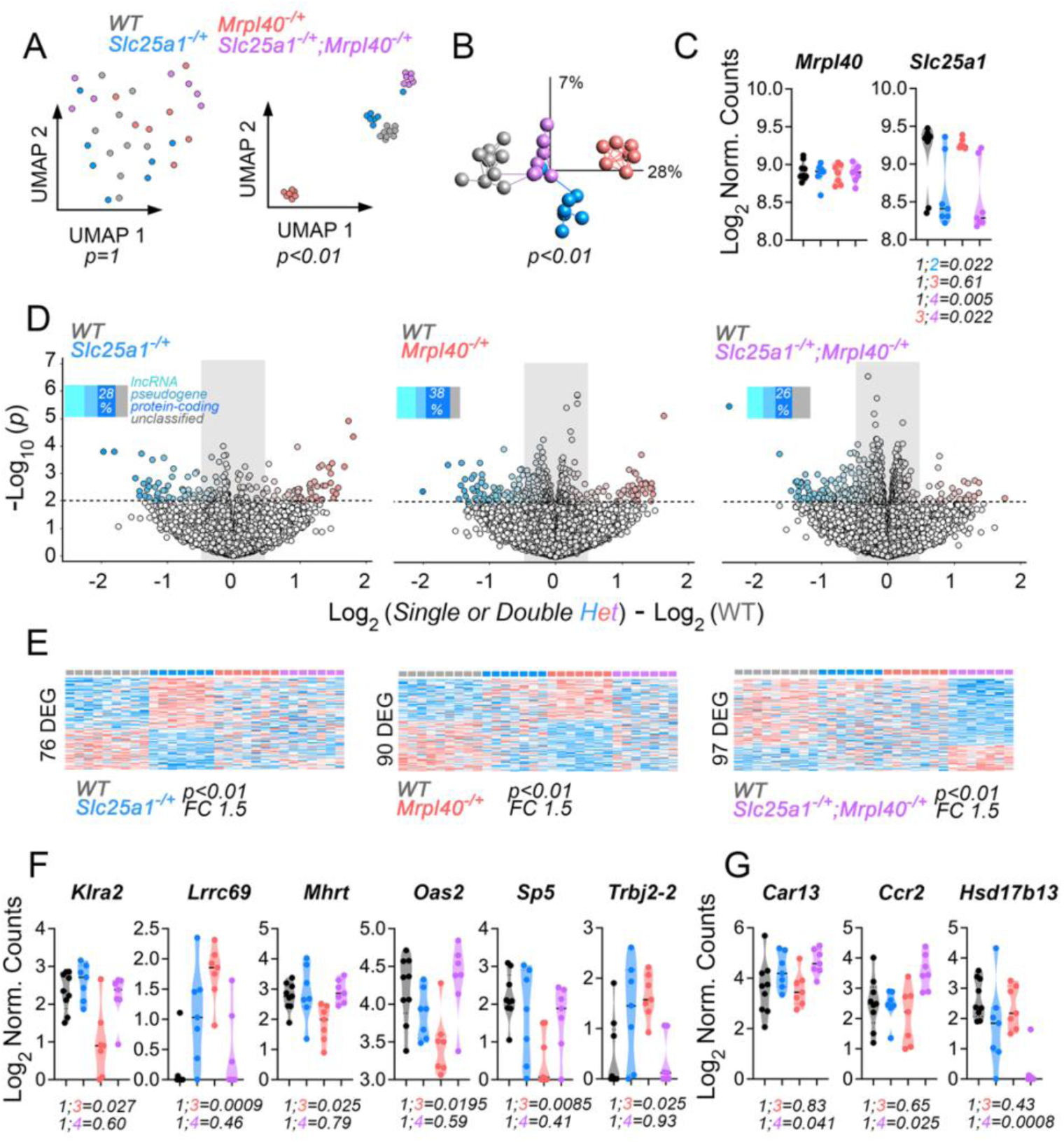
Cortical Transcriptome of Single and Double *Slc25a1^-/+^*;*Mrpl40^-/+^* Heterozygotic Mice. **A**. Uniform Manifold Approximation and Projection (UMAP) of cortical transcriptome before and after one-way ANOVA thresholding by p<0.01 of the indicated genotype transcriptomes. **B**. PCA of transcriptomes after one-way ANOVA thresholding by p<0.01. Lines represent Euclidean distance clustering. **C.** Normalized mRNA counts for Mrpl40 and Slc25a1 transcripts. Kruskal-Wallis test followed by Benjamini and Hochberg multiple corrections. **D**. Volcano plots and **E**. heat maps of paired comparisons between wild type animals with single or transheterozygotic animals. Insets in volcano plots show percentage of diverse types of RNAs differentially expressed. Heat Maps depict z-scored hierarchical clustering of transcripts after thresholding with a cut-off fold of change=1.5 and p<0.01 (Welsh t test). **F** and **G** depict violin plots of mRNAs either rescued (F) or worsened (G) in *Slc25a1^-/+^*;*Mrpl40^-/+^*cortex. Kruskal-Wallis test followed by Benjamini and Hochberg multiple corrections. All panels wild type n=9 and all other genotypes n=7.

## Discussion

A frequent strategy to assess the contribution of genes to copy number variation syndrome phenotypes is the identification of driver genes whose haploinsufficiency is sufficient to elicit phenotypes. One example of this is Tbx1 in the 22q11.2 microdeletion. Tbx1 haploinsufficiency causes conotruncal cardiovascular, cranial, and brain development, as well as behavioral phenotypes observed in 22q11.2 microdeletion syndrome models ^43, 45–47^. We previously expanded the repertoire of 22q11.2 genes required for cardiac development to include *Slc25a1* ^30^. Our findings add *Mrpl40* to the list of haploinsufficiency genes affecting brain and cardiac phenotypes, supporting the concept that multiple loci within 22q11.2 determine tissue phenotypes. The focus on single driver genes, while an important approach, neglects the possibility that multiple genes within a microdeleted chromosomal locus could interact to specify tissue-specific phenotypes. Human genomic studies and direct hypothesis testing in *Drosophila* show that rather than driver genes specifying phenotypes, traits are determined by gene interactions that enhance or ameliorate a phenotype. This idea has been elegantly demonstrated in *Drosophila* where tissue phenotypes are modulated by interactions between pairs of genes encoded within a human CNV locus ^10, 11, 13, 14, 48^. Here we provide evidence of synthetic interactions between two of the eight mitochondrial genes located in the 22q11.2 microdeletion locus. We demonstrate that *Mrpl40* and *Slc25a1* genetically interact to specify cardiac development, behavior, and brain transcriptomes. Even though the two genes form part of a biochemical mitochondrial interactome and are both required for normal mitochondrial function, the combined haploinsufficiency of these genes is suppressive at the level of tissue phenotypes. These results reveal that the robustness of the principles governing genetic interactions among CNV genes discovered in *Drosophila* also apply to mammals.

The suppressive outcome in heart and brain of *Slc25a1^-/+^*;*Mrpl40^-/+^* mice is surprising. Our biochemical studies show that proteins belonging to the SLC25A1 interactome are affected by knocking out MRPL40, thus revealing possible mitochondrial convergence points between SLC25A1 and MRPL40. Such convergence points include the function of the respiratory chain, the mitochondrial ribosome, the maturation of the mitochondrial transcriptome, possible activation of innate immune responses by excess mitochondrial nucleic acids, alterations in acetyl-CoA pools, and cellular processes that require this metabolite, such as epigenome modifications. These convergence points could result in synthetic interactions enhancing or ameliorating a phenotype. There is precedent for suppressive interactions even within components of the respiratory chain. For example, mutations in complex I subunits can be suppressed by intra complex I mutants ^49^. Presently, we do not know which one of these MRPL40- and SLC25A1-dependent mitochondrial mechanisms could account for suppressive effects. However, we speculate about three possibilities. Suppressive interactions may be related to pools of unassembled mitochondrial ribosome proteins, either within mitochondria or in the cytoplasm. We have shown that SLC25A1 KO cells have decreased integrity of the mitochondrial ribosome and reduction of the levels of mitochondrial ribosome proteins ^29^. If phenotypes observed in tissues are a consequence of unassembled mitochondrial ribosome proteins acting as dominant negative molecules, then reducing their levels would prevent such effect. Decreasing MRPL40 gene dosage could reduce the levels of unassembled mitochondrial ribosome proteins in an SLC25A1-deficient background, thus preventing a dominant effect. A second putative mechanism is heterogenous cell responses to *Mrpl40* haploinsufficiencies. The levels of mitochondrial ribosome mRNAs are cell type specific, with interneurons expressing the highest levels of these transcripts ^29, 50^. Cell types with higher levels of MRPL40 expression could potentially have a greater suppressive effect due to greater decrease in mitochondrial ribosome gene expression. The regional specificity of the gene expression changes in the brain comparing cortex and hippocampus between *Slc25a1^-/+^* and *Mrpl40^-/+^* supports this idea. Finally, a third mechanism is impaired handling of mitochondrial or cytoplasmic calcium. A prior *Mrpl40^-/+^* mouse model shows increased mitochondrial calcium transients and short-term potentiation, the latter a phenotype that could be modulated by the expression of the mitochondrial ADP-ATP translocator Slc25a4. Slc25a1 and Slc25a4 biochemically interact and their knockouts have decreased calcium uptake into mitochondria, thus disruption of *Slc25a1* could restore mitochondrial calcium homeostasis in *Mrpl40^-/+^*neurons ^22, 51^. We focused on genetic interactions between two of six mitochondrial genes encoded in the 22q11.2 microdeleted locus. However, defects in the 22q11.2 mitochondrial genes Txnrd2, Prodha, Snap29 and Comt cause defective neurodevelopment and/or behavioral alterations ^52–58^. Therefore, we postulate further synthetic interactions exist between additional 22q11.2 mitochondrial genes, which may be difficult to predict from studies in single gene heterozygosity models. Most neurodevelopmental syndromes are now understood to have a polygenic basis, which provides rationale for expanded interrogation of CNVs and multigenic models ^11, 12, 59^.

We found defective prepulse inhibition in our *Mrpl40^-/+^* animals, a phenotype that aligns with prior results in *Danio rerio mrpl40^-/-^* displaying altered acoustic startle responses, and with mutations in the mitochondrial genome gene *MT-TL1*, which encodes a tRNA required for protein synthesis in mitochondria ^53, 60^. Heteroplasmic mice carrying this mitochondrial genome mutation also have reduced prepulse inhibition ^60^. Our results stand in contrast to a previous study in another *Mrpl40^-/+^*mouse model, which reported normal prepulse inhibition ^51^. These mice share similar genetic backgrounds in these two *Mrpl40^-/+^* mice (MGI:3847513 and MGI:4431658), so we suspect the discrepancy in prepulse inhibition across these two models is the age at which tests were performed. Age is a variable with strong effects in prepulse inhibition, particularly in the C57BL/6N strain used in our studies ^61–63^. We tested our animals at 9-11 weeks of age, while Devaraju et al. tested them between 16-20 weeks. As further confirmation, we demonstrated that the magnitude of prepulse inhibition reduction in *Mrpl40^-/+^* mice is similar to that in Df1/+ mice, a 22q11.2 microdeletion animal model, which we tested with the same behavioral experimental paradigm and at the same age. In summary, our *Mrpl40* mouse model adds to the list of genes within the 22q11.2 locus that impair prepulse inhibition. Our findings also suggest a more complex genetic regulation of prepulse inhibition responses among the genes affected in the 22q11.2 locus. Multiple CNVs of the mouse 22q11.2 synthenic region show defective prepulse inhibition, including Df1/+, Df(16)A+/−, LgDel/+, Df(h22q11)/+, Del(1.5 Mb)/+, and Del(3.0 Mb)/+ ^42, 64–69^. The genetic defect in four of these mouse models spans *Slc25a1* and *Mrpl40,* yet they still show robust reduction of prepulse inhibition (Df(16)A+/−, LgDel/+, Del(1.5 Mb)/+, Del(3.0 Mb)/+). If *Slc25a1^-/+^* can suppress a *Mrpl40^-/+^* prepulse phenotype, as we have shown here, then other genes within the 22q11.2 locus and their interactions must be causal of a prepulse inhibition defects. Among these genes, haploinsufficiency of either Tbx1 ^44, 70, 71^, or the mitochondrial gene Prodh ^56, 58^ cause this phenotype.

A limitation of our studies is the correlative nature of the transcriptome and behavioral phenotypes, as well as the fact that our transcriptomes do not include brain regions engaged in prepulse inhibition responses, including the amygdala and midbrain ^72^. *Slc25a1* and *Mrpl40* are broadly expressed genes; thus, a reasonable assumption is that the genetic interaction we identify occurs in a cell-autonomous and anatomically localized manner. However, it is possible that the suppressive effect between these haploinsufficient genes could be distributed across the brain regions participating in prepulse inhibition. *Slc25a1* and *Mrpl40* cell-autonomous suppressive interactions are more likely in heart than brain as a result of the heart’s more homogenous cellular and anatomical landscape. Our emphasis has been in the most salient phenotypes caused by *Slc25a1* and *Mrpl40* haploinsufficiency and their suppressive interactions. However, even though brain transcriptomes show discrete genes whose expression is restored to wild type levels in combined heterozygotic tissue, the paucity of altered mRNAs prevents the identification of potential cell autonomous molecular mechanisms affected by the single and combined heterozygosity.

Together, our findings argue that the genetic architecture of the 22q11.2 microdeletion emerges from combinatorial interactions among multiple loci, including mitochondrial genes within the CNV. By uncovering a surprising suppressive interaction between *Slc25a1* and *Mrpl40*, we show that mitochondrial gene dosage can buffer or reshape phenotypes *in vivo*, influencing cardiac development, brain transcriptomes, and sensorimotor gating. The presence of prepulse inhibition deficits in *Mrpl40^-/+^* mice, their alignment with mitochondrial ribosome and mitochondrial-genome mutants, and their persistence across multiple 22q11.2 CNV models underscore both the centrality of mitochondrial pathways to neural circuit function and the likelihood that additional 22q11.2 genes participate in genetically interacting modules that shape this behavioral outcome. Although the precise mitochondrial mechanisms driving suppression in brain and heart remain unresolved, our results highlight the complexity of mitochondrial genetic networks operating during development. Ultimately, this work advances a network-based model of CNV pathology in which interacting mitochondrial and non-mitochondrial genes collectively sculpt tissue phenotypes and modulate their penetrance.

## Materials and Methods

### Cell Culture and MRPL40 KO Cells Generation

Crispr edited SH-SY5Y cells were generated at Synthego with guide sequence UCACCCGCUAGUCGGGCGCA. Standard SH-SY5Y growth media was further supplemented with uridine (50 µg/mL), sodium pyruvate (110 µg/mL), and D-glucose (2 mg/mL). MRPL40 knock-out (KO) and control SH-SY5Y cell pools were cloned in-house. To verify isolated clones, DNA was extracted using QuickExtract DNA extraction solution (Lucigen QE0905T) according to manufactures protocol. The MRPL40 targeted genetic region was amplified with forward (GCAGCTGACACCCTAGGC) and reverse (TTCTCTCCCACTTCACAGGAAAAT) primers using AmplitaqGold 360 Master Mix (Life Technologies 4398876). PCR products were Sanger sequenced and Inference of CRISPR Edits (ICE) analysis (Synthego.com) was used to confirm genetic mutation. Western blot was used to confirm loss of MRPL40 protein.

All cells were maintained in DMEM supplemented with 10% FBS, and non-essential amino acids (NEAA) and uridine, D-glucose, and sodium pyruvate as described above. Control and MRPL40 KO clones were cultured in the supplemented media, in a humidified incubator at 37°C and 5% CO_2_.

To generate stable cell lines, SH-SY5Y cells (ATCC, CRL-2266; RRID:CVCL_0019) were transfected with ORF expression clone containing C terminally tagged Myc-DDK MRPL40 (Origene, RC202166) or N terminally tagged FLAG-SLC25A1 (GeneCopoeia, EX-A1932-Lv1020GS) as described ^29^. The stable cell lines were maintained in DMEM media containing 10% FBS, 100 µg/ml penicillin and streptomycin, and neomycin 0.2 mg/ml (Hyclone, SV30068) or puromycin 2 µg/ml respectively (Invitrogen, A1113803) at 37°C in 10% CO_2_

### Antibodies and Immunoblots

For immunoblot, cells were washed twice with phosphate buffered saline (PBS) supplemented with 1 mM MgCl_2_ and 100 µM CaCl_2_, then lysed with Buffer A (10 mM HEPES pH 7.4, 150 mM NaCl, 1 mM EGTA, and 0.1 mM MgCl_2_, with 0.5% Triton X-100 and Complete anti-protease) Lysates were incubated at 4°C for 30 minutes with periodic vortexing and clarified by centrifugation at 20,000 RCF for 15 minutes. The supernatant was collected for downstream use and the pellet was discarded. Clarified cell lysis was diluted to 1 µg/µL and Laemmli buffer added for SDS-PAGE. Equivalent amounts of sample were loaded onto 4-20% Criterion TGX Midi-gels for SDS-PAGE and transferred to PVDF at 1.5 mA/cm^2^ for 65 minutes using a semi-dry method. Membranes were blocked for 30 minutes at room temperature with tris-buffered saline with 0.05% Triton-X 100 (TBST) containing 5% non-fat milk, washed well, and incubated overnight at 4°C with primary antibody diluted in antibody base (3% BSA and 0.2% sodium azide in PBS). The following day, membranes were washed in TBST and incubated with secondary HRP conjugated antibody or rhodamine conjugated anti-tubulin diluted in TBST with 5% non-fat dry milk, for 30 minutes at room temperature. Washed membranes were imaged either with ChemiDoc MP (BioRad) fluorescent detection or with Western Lighting Plus ECL substrate incubation (PerkinElmer, NEL105001EA) for chemiluminescent detection on GE Healthcare Hyperfilm. Quantification of western blot band intensity was determined with Fiji (ImageJ) or Image Lab (BioRad). The intensity of the KO band was divided by the average intensity of the WT bands on the membrane for that particular antibody.

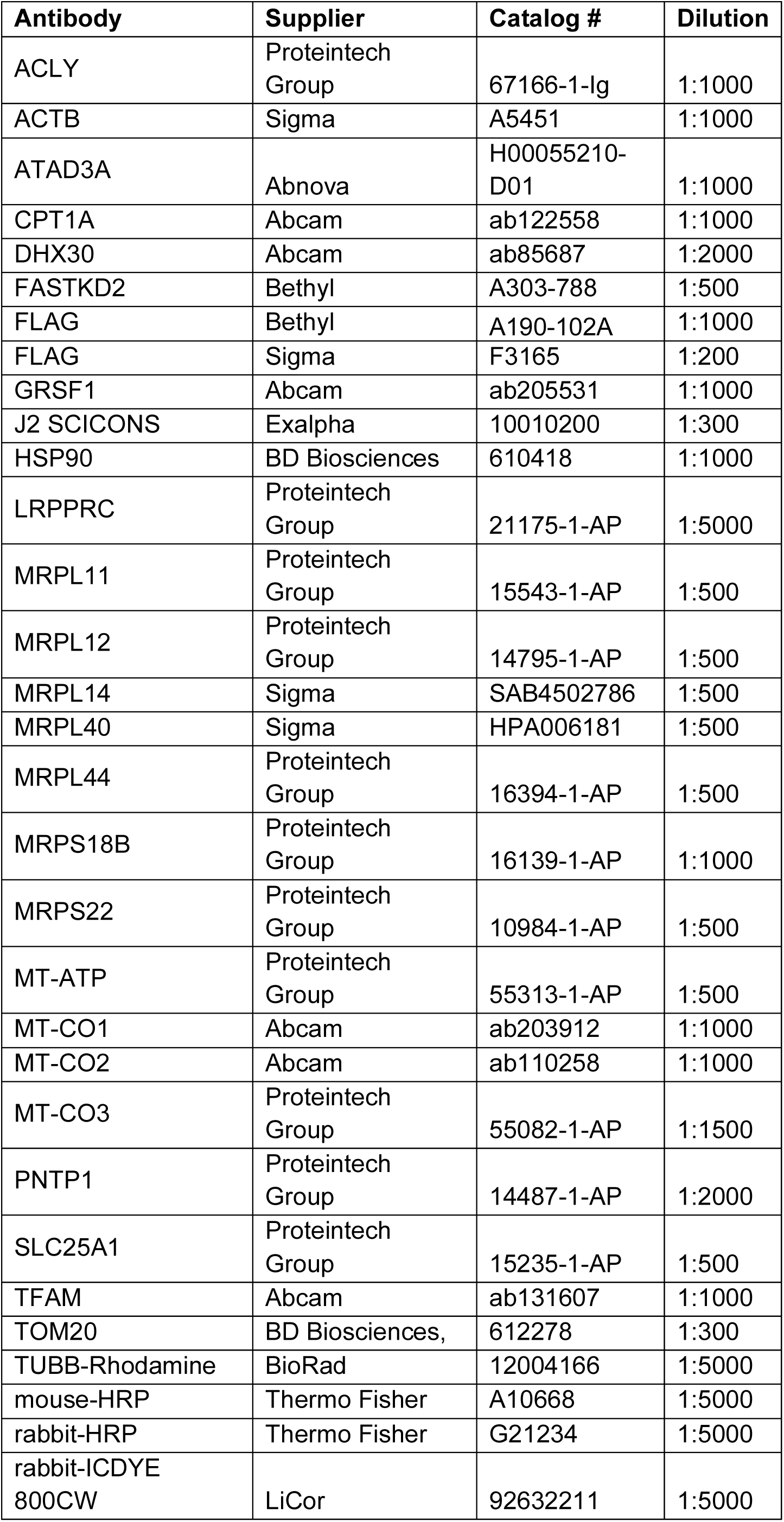

| **Antibody** | **Supplier** | **Catalog #** | **Dilution** |
| --- | --- | --- | --- |
| ACLY | Proteintech Group | 67166-1-Ig | 1:1000 |
| ACTB | Sigma | A5451 | 1:1000 |
| ATAD3A | Abnova | H00055210-D01 | 1:1000 |
| CPT1A | Abcam | ab122558 | 1:1000 |
| DHX30 | Abcam | ab85687 | 1:2000 |
| FASTKD2 | Bethyl | A303-788 | 1:500 |
| FLAG | Bethyl | A190-102A | 1:1000 |
| FLAG | Sigma | F3165 | 1:200 |
| GRSF1 | Abcam | ab205531 | 1:1000 |
| J2 SCICONS | Exalpha | 10010200 | 1:300 |
| HSP90 | BD Biosciences | 610418 | 1:1000 |
| LRPPRC | Proteintech Group | 21175-1-AP | 1:5000 |
| MRPL11 | Proteintech Group | 15543-1-AP | 1:500 |
| MRPL12 | Proteintech Group | 14795-1-AP | 1:500 |
| MRPL14 | Sigma | SAB4502786 | 1:500 |
| MRPL40 | Sigma | HPA006181 | 1:500 |
| MRPL44 | Proteintech Group | 16394-1-AP | 1:500 |
| MRPS18B | Proteintech Group | 16139-1-AP | 1:1000 |
| MRPS22 | Proteintech Group | 10984-1-AP | 1:500 |
| MT-ATP | Proteintech Group | 55313-1-AP | 1:500 |
| MT-CO1 | Abcam | ab203912 | 1:1000 |
| MT-CO2 | Abcam | ab110258 | 1:1000 |
| MT-CO3 | Proteintech Group | 55082-1-AP | 1:1500 |
| PNTP1 | Proteintech Group | 14487-1-AP | 1:2000 |
| SLC25A1 | Proteintech Group | 15235-1-AP | 1:500 |
| TFAM | Abcam | ab131607 | 1:1000 |
| TOM20 | BD Biosciences, | 612278 | 1:300 |
| TUBB-Rhodamine | BioRad | 12004166 | 1:5000 |
| mouse-HRP | Thermo Fisher | A10668 | 1:5000 |
| rabbit-HRP | Thermo Fisher | G21234 | 1:5000 |
| rabbit-ICDYE<br>800CW | LiCor | 92632211 | 1:5000 |

### Immunoprecipitation Assays

Immunoprecipitation experiments were performed as described in Gokhale et. Al 2021. Briefly, SH-SY5Y cells expressing either FLAG-SLC25A1 or Myc-DDK MRPL40 were grown in 10 cm tissue culture dishes, placed on ice, and rinsed twice with cold PBS (Corning, 21-040-CV) containing 0.1 mm CaCl_2_ and 1.0 mm MgCl_2_. Cells were lysed in a buffer containing 150 mm NaCl, 10 mm HEPES, 1 mm EGTA, and 0.1 mm MgCl_2_, pH 7.4 (Buffer A) with 0.5% Triton X-100 and Complete anti-protease (Roche, 11245200). The lysates were placed on ice for 30 min and centrifuged at 16,100 × *g* for 10 min. The clarified supernatant was retrieved, and protein concentration was determined using the Bradford Assay (Bio-Rad, 5000006). For immunoprecipitation assays, 500 µg of the soluble protein lysate was applied to 30 µl Dynal magnetic beads (Invitrogen, 110.31) coated with 1 µg of the mouse monoclonal FLAG antibody. This mixture was then incubated on an end-to-end rotator for 2 h at 4°C. As controls, the immunoprecipitation was outcompeted with the 3XFLAG peptide (340 μm; Sigma, F4799). After 2 hours, the magnetic beads were washed 6 times with Buffer A with 0.1% Triton X-100. Proteins were eluted from the beads with Laemmli buffer. Samples were then analyzed by immunoblot.

### Seahorse

SH-SY5Y control and MRPL40 KO clones were plated at 32,000 cells per well in an Agilent Seahorse culture plate (Agilent, Seahorse FluxPak Kit 103792-100) with DMEM supplemented with 10% FBS, uridine, additional D-glucose, sodium pyruvate and NEAA and incubated overnight at 37°C, with 5% CO_2_. The flux plate cartridge was hydrated with calibrant solution overnight at 37°C, without injected CO_2_. The following day, culture media was removed, washed twice with Seahorse assay media (103574-100), adjusted to a final volume of 180 µL per well, then incubated for 1 hour before the start of the mitochondrial stress test, at 37°C without injected CO_2_. Mitochondrial stress test drugs were loaded into injection ports at a 10x concentration to yield final concentrations of 2 µM oligomycin, 0.25 µM FCCP, 1 µM rotenone, and 1 µM antimycin A. The standard Agilent mitochondrial stress test protocol was used, including three baseline measurements prior to injection and three measurements following each injection of oligomycin, FCCP, and rotenone/antimycin A. Agilent WAVE software was used to measure and analyze oxygen consumption rates (OCR) and extracellular acidification rates (ECAR). Data were normalized by total protein content per as determined by BCA assay. Briefly, cells were washed with PBS supplemented with 1 mM MgCl_2_ and 100 µM CaCl_2_, then lysed in-well with 0.5% Triton X-100/Buffer A (as described above). BCA reagent was added to the lysis in each well and incubated at 37°C for 30 minutes before reading on the BioTech plate reader at 562 nm against BSA standards.

### qRT-PCR

Total RNA was extracted from roughly 6.5 million control and MRPL40 KO SH-SY5Y cells with Trizol reagent, according to manufacturer’s protocol. First strand cDNA synthesis was done with the SuperScript III Kit (Thermo Fisher 12574030) with random hexamers and 5 µg of total RNA following manufactures protocol. RT-PCR of the resulting cDNA was performed in triplicate on the QuantStudio 6 Flex (Applied Biosystems) with LightCycler 489 SYBR Green I Master Mix (Roche 04707516001) and primers for MTRNR1, MTRNR2 and Vimentin as ^29^. Data analysis was completed using QuantStudio RT-PCR software version 1.2, with relative quantification based on standard curves generated from control SH-SY5Y cells.

### Mitochondrial Genome Quantifications

Total DNA was extracted from approximately 2.5 million control and MRPL40 KO SH-SY5Y cells using the DNeasy Blood and Tissue DNA Extraction Kit (Qiagen 69504) according to manufactures guidelines. DNA concentration and purity were validated on the Nanodrop OneC. Following the protocol of Rooney et al. for determination of mitochondrial DNA copy number, 6 ng of total DNA was analyzed on the QuantStudio 6 Flex (Applied Biosystems) with LightCycler 489 SYBR Green I Master Mix (Roche 04707516001) and primers for mitochondrial specific (MT-tRNA-Leu) or nuclear specific (Nuc-Beta2-microglobulin) DNA. Ct values determined by QuantStudio RT-PCR software were used to calculate the relative mitochondrial DNA as shown by ^73^.

1. (ΔCt) = (Nuclear DNA C_t_) – (Mitochondrial DNA C_t_)
2. Relative mitochondrial DNA = 2 × 2^ΔCT^

### Nanostring Determinations

Cells were solubilized in TRIzol (Invitrogen). RNA isolation and NanoString analysis was performed by the Emory Integrated Genomics Core. RNA quality was assessed by bioanalyzer. mRNA counts were normalized to the housekeeping gene CLCT for the MitoString using nSolver and then processed with Qlucore Omics Explorer Version 3.6 Software.

### Confocal Microscopy

Cells were fixed in 4% PFA at 37°C for 10 minutes and permeabilized with 0.25% Triton X 100 for 5 minutes with 20U/mL SUPERase-In RNAse inhibitor. Cells were blocked with 10% fetal bovine serum in PBS and incubated with primary antibodies: J2 and TOM20 (BD Biosciences, 612278). Slides were imaged with a Nikon A14 HD25 confocal microscope and 60x Apo lens (NA=1.4, WD=140um). dsRNA localized to the mitochondria (TOM20) was quantified with ImageJ/Fiji software in the focal plane of maximal intensity. dsRNA was measured as pixel intensity per cell, restricted to TOM20 segmented by thresholding.

### Animals

All animal procedures were approved by Emory University’s Institutional Animal Care and Use Committee. SLC25A1 KO mice (C57BL/6N-*Slc25a1tm1a(EUCOMM)Wtsi*/Mmucd) were purchased from the Mutant Mouse Resource and Research Centers (MMRRC 042258-UCD) and genotyped according to the MMRRC PCR protocol. The WT allele was detected by primers GAATTGGTCGTGGTCTCAGTAGCC and GGAGTGCCCAAGAGACTCTGAGC producing a 366 base pair product. The KO allele recognized primers GAGATGGCGCAACGCAATTAATG and TAGTGAGTTATGCTTTGAAGACTTCGC producing a 253 base pair product.

DF1 mice (B6.129S7-Del(16Es2el-Ufd1l)217Bld/Cnrm) were from Adriano Buzzati-Traverso at the Institute of Genetics and Biophysics, National Research Council through the European Mouse Mutant Archive (EMMA) managed by Intrafrontier (Intrafrontier EM02122). Genotyping was performed with primers TGGGCAATTGTTTAATCTTCC and TCTTTGTCAGCAGTTCCCTTT, where the mutant gene generated a 290 base pair product and the wild type gene provided a 150 base pair product.

We used the CRISPR/Cas9 system to generate the Mrpl40 knockout mouse line. One 20 base pair Mrpl40 guide RNA (gRNA) sequence, ACCTTCTTCTTCTTCCGCA, was designed at the syntenic loci in the mouse genome. The Emory Mouse Transgenic and Gene Targeting core injected 20 ng/µL of gRNA and 20 ng/µL of Cas9 protein into single-cell C57BL/6J zygotes. Embryos were cultured overnight and transferred to pseudopregnant females. The resulting pups were screened for Mrpl40 knockout via PCR. To confirm the desired mutation, Sanger sequencing was performed on purified PCR DNA from the potential mutant mice. Heterozygous mice progress to maturity and produce viable pups. Genotyped was done with primers CACTTGTTCCTACCACAGACATG and GACAGTGGACTAAGCTCGTGGAG. The resulting PCR product from the WT allele was digestible by AciI, producing bands of 237 and 136 base pairs. However, the MRPL40 mutant allele product was undigestible and produced a PCR product 373 base pairs in length.

### Heart Histology and Analyses

Timed matings were conducted using *Slc25a1^+/-^* male and female mice aged 2–3-months, with the detection of a copulation plug designated as embryonic day (E) 0.5. Embryos were harvested at E14.5, fixed in 10% formalin and embedded in paraffin. Paraffin-embedded embryos were sectioned at 6 µm, followed by deparaffinization, rehydration, and staining with hematoxylin and eosin (H&E). Slide were imaged using a Nanozoomer 2.0-HT whole-slide scanner (Hamamatsu). Myocardial trabecular and compact layer thicknesses was quantified on H&E-stained sections using QuPath software ^74^.

### Behavioral Analyses

#### Morris Water Maze

Morris Water Maze training took place in a round, water-filled tub (52 inch diameter) in an environment rich with extra maze cues and a small platform 1 cm below the surface (see below). White tempera paint (non-toxic) was added to the water to make it opaque so that the mice could not see the platform. Mice were placed in the water maze with their paws touching the wall from 4 different starting positions (N,S,E,W) in water that started at 25°C and typically declined to 22°C by the time a whole group of mice was tested. An invisible escape platform was in the same spatial location 1 cm below the water surface independent of a subjects start position on a particular trial. In this manner subjects were able to utilize extra maze cues to determine the platform’s location. Each subject was given 4 trials/day for 5 days with a 15-min inter-trial interval. The maximum trial length was 60 s and if subjects did not reach the platform in the allotted time, they were manually guided to it. Upon reaching the invisible escape platform, subjects were left on it for an additional 5 s to allow for survey of the spatial cues in the environment to guide future navigation to the platform. After each trial, subjects were dried and kept in a dry plastic holding cage filled with paper towels to allow them time to dry off. The holding cage was placed half-on, half-off a heating pad and mice were monitored closely. Following the 5 days of task acquisition, a probe trial was presented during which time the platform was removed and the amount of time and distance swam in the quadrant which previously contained the escape platform during task acquisition was measured over 60 s. All trials were videotaped and performance analyzed by means of MazeScan (Clever Sys, Inc.).

#### Elevated Plus Maze

The elevated plus maze, which is constructed of Plexiglas, consisted of two open arms and two enclosed arms arranged in a plus orientation. The arms were elevated 30 inches above the floor, with each arm projecting 12 inches from the center. To begin each test, mice were placed in the center of the maze facing one of the open arms and allowed to freely explore the apparatus for five minutes, during which time their behavior were videotaped. Mice were returned to their home cage at the end of the 5 min test.

#### Prepulse Inhibition (PPI)

Prepulse inhibition was assessed in a sound attenuated chamber (San Diego Instruments). In the PPI test mice following a 5 minute habituation period, subjects were presented with 66 total trials in a pseudorandom order. Those trials consisted of startle stimulus alone trials (120 dB, 20 ms), prepulse alone trials (20 ms white noise at 70, 74, 78, 82, or 86 dB), and prepulse and startle stimulus combined trials (each of the 5 prepulse intensities followed by a 20 ms 120-dB startle stimulus). Each of the trial types was presented in a pseudorandom fashion such that each trial was presented 6 times and no two consecutive trials were identical. Mouse movement was measured by a piezoelectric accelerometer during 100 ms after startle stimulus onset for 100 ms. PPI (%) was calculated. At the end of the session, subjects were removed from the sound attenuated chamber and placed in their home cage.

#### Circadian Activity

Mice were placed in plexiglass activity cages (with appropriate clean bedding) equipped with infrared photobeams (San Diego Instruments) for 23 hours. Ambulations (consecutive beam breaks) were counted by a computer. Food and water will be available ad libitum.

#### Marble Burying

A standard mouse cage lined with corncob bedding was filled with 20 glass marbles (15 mm diameter), arranged in a 4 x 5 matrix and equidistant from one another. Mice were then placed in the clean cage with the marbles for 30 min. The number of marbles buried (>50% marble covered by bedding) was recorded as the primary dependent variable

#### Nestlet Shredding

The mice were moved from their home cages to individual testing cages. Each testing cage contained 3.0 g of Nestlets (commercially available pressed cotton squares), which mice typically use to make nests. The cages did contain any other environmental enrichment items. Mice were left in the testing cages for 120 mins. At the end of the 120 mins, unused nest material was collected and weighed.

### RNAseq

RNA Isolation. Admera Health performed all RNA isolation, library preparation, and sequencing. Qiazol phase separation, followed by cleanup with RNeasy 96 was used to isolate RNA from Hippocampal and Cortical samples. Both RNA Tapestation assay and High Sensitivity RNA Tapestation assay (Agilent Technologies Inc., California, USA) and quantified by Infinite F Nano+ 200 Pro Tecan (Tecan, Switzerland) assessed the quality of the isolated RNA.

Coding RNA transcripts were isolated using NEBNext® Poly(A) mRNA Magnetic Isolation beads as part of the NEBNext® Ultra™ II Directional RNA Library Prep Kit for Illumina® (New England BioLabs Inc., Massachusetts, USA). cDNA synthesis and library preparation was performed using the same kit. Prior to first strand synthesis, samples are randomly primed (5’ d(N6) 3’[N=A,C,G,T]) and fragmented. The Protoscript II Reverse Transcriptase with a longer extension period, approximately 30 minutes at 42⁰C synthesized the first strand of cDNA. Final library quantification was performed by Qubit 2.0 (ThermoFisher, Massachusetts, USA) and quality assessment was made by TapeStation HSD1000 ScreenTape (Agilent Technologies Inc., California, USA). Final library size was about 450bp with an insert size of about 300bp. Illumina® 8-nt dual-indices were used. Equimolar pooling of libraries was performed based on QC values and sequenced on an Illumina NovaSeq X Plus (Illumina, California, USA) with a read length configuration of 150 PE for 60M PE reads per sample (30M in each direction).

RNA sequencing analysis: FastQC removed samples of poor quality^75^). Trimmomatic (version v0.39) removed and trimmed adapter sequences and readings of poor quality. The web interface and public servers, usegalaxy.org and usegalaxy.eu, was used for mapping and analysis of RNAseq reads ^76^. The Galaxy server running Hisat2 (Galaxy Version 2.2.1+galaxy0), FeatureCounts (Galaxy Version 1.6.4), and Deseq2 (Galaxy Version 2.11.40.8+galaxy1) was used to map sequence reads^77–79^. FeatureCounts files and raw files are available at GEO with accession XXX. Hisat2 was run with the following settings: paired-end, stranded, default settings (except for when a GTF file was used for transcript assembly). For GTF files, we used the Mus musculus (Mouse), Ensembl, GRCm39 build from iGenome (Illumina). The aligned SAM/BAM files were processed using Featurecounts with Default settings, except we used the Ensembl GRCm39 GTF file and output for DESeq2 and a gene length file.

Gene counts were normalized using DESeq2 (Love et al., 2014) followed by a regularized log transformation. Differential Expression was determined by DESeq2; the factors used were tissue type (hippocampus and cortex), Mrpl40 expression (+/+ and -/+), and Slc25a1 expression (+/+ and -/+). Pairwise comparisons were done across Mrpl40 and Slc25a overexpression status. Animal genotypes: Mrpl40 +/+ Slc25a1 +/+; Mrpl40 -/+ Slc25a1 +/+; Mrpl40 +/+ Slc25a1 -/+; Mrpl40 -/+ Slc25a1 -/+. All normalized tables, including regularized log transformation and variance stabilized tables, were generated using size estimation, where the standard median ratio was used; the fit type was parametric, and outliers were filtered using a Cook’s distance cutoff.

### Data Access

RNAseq data are accessible through the GEO portal GSE315447

### Statistics

ANOVA analyses were conducted with Prism Version 10.2.2. Data were tested for normality and log transformed when required for ANOVA analyses. Two-sided permutation t-test were done using the estimationstats engine ^80^ and Kolmogorov-Smirnov tests were conducted with the engine http://www.physics.csbsju.edu/stats/KS-test.n.plot_form.html.

Coessentiality Analysis was performed with the FIREWORKS engine ^34^ using SLC25A1 and MRPL40 genes as entries in the Pan-Cancer dataset as Context. The engine was run with the top 10 primary nodes and the top five secondary nodes positively and negatively correlated. Data were exported and analyzed in Cytoscape 3.10.2. Nodes enriched annotations were inspected with ENRICHR ^81^.

## Acknowledgements

This work was supported by NIH grants 1RF1AG060285 to VF, R01ES034796 to EW and AG, and K01MH133970 to RHP. JQK Additional Ventures Single Ventricle Research Fund, the Department of Defense (0000063651), and the NIH R01GM144729. HM-V is supported by an ARCS Foundation Award, John B. Lyon Memorial Scholarship Award, and HHMI Gilliam Award. This study was supported in part by the Emory Transgenic Mouse and Genomics Cores, which are subsidized by the Emory University School of Medicine. VF is grateful for mitochondria provided by Maria Olga Gonzalez.

## Supplementary Figures, Legends, and Tables

**Supplementary Fig. 1.**
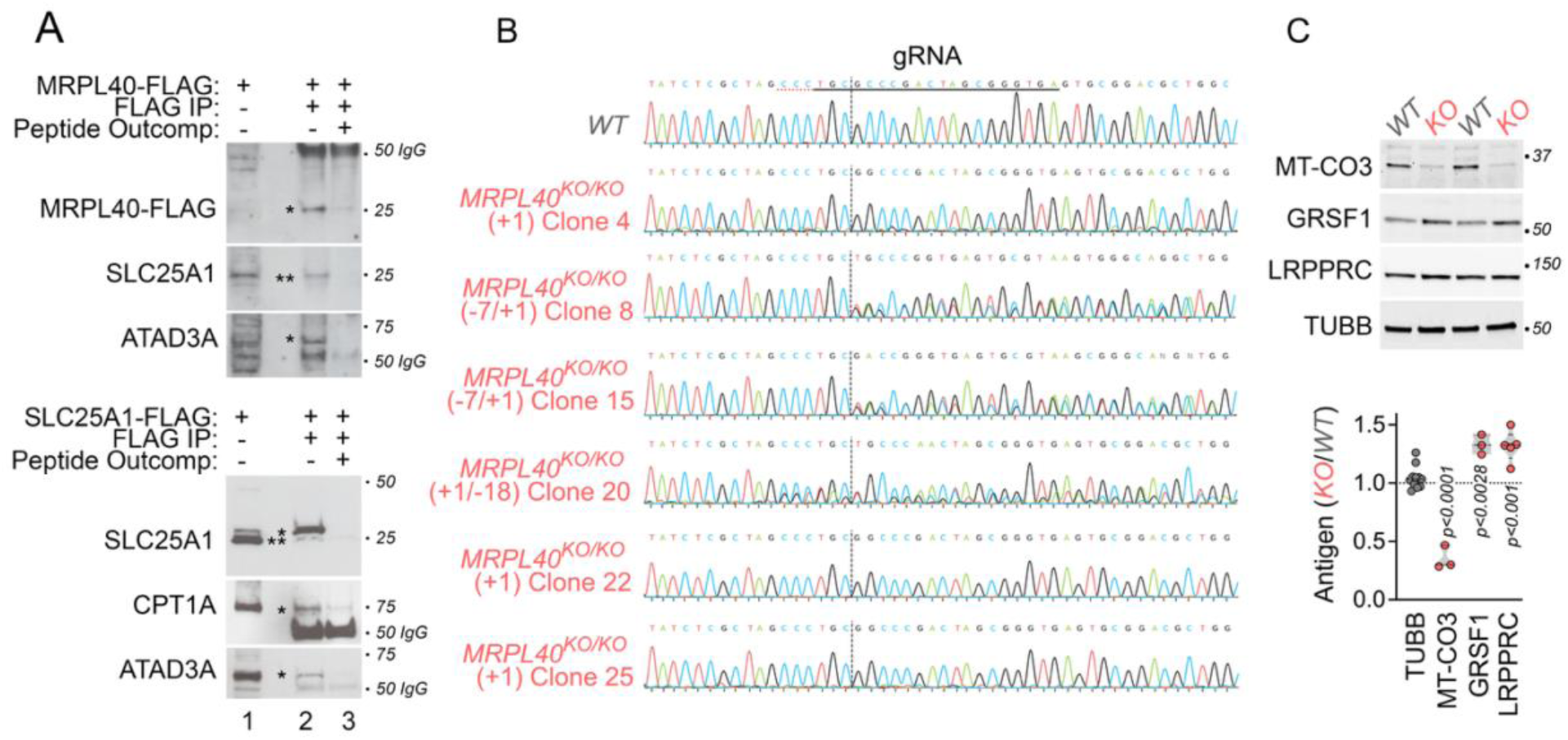
MRPL40 and SLC25A1 Coprecipitate plus CRISPR MRPL40 Null Mutants Sequences and Additional Characterizations. **A.** MRPL40 and SLC25A1 coprecipitate. Extracts from neuroblastoma cells stably expressing either MRPL40-FLAG or SLC25A1-FLAG were immunoprecipitated with FLAG antibodies (lanes 2-3) in the absence or presence of an excess FLAG peptide (lane 3). Lane 1 corresponds to input. Asterisks mark antigens tested. Double asterisk marks endogenous SLC25A1 and single asterisk labels SLC25A1-FLAG. **B.** Sanger sequencing chromatograms of wild type and MRPL40 null clones (See Fig. 1B). Homozygotes and compound heterozygote clones are presented. Clone 8 has a deletion of 7bp and 1 bp insertion (-7/+1). **C**. Immunoblot of the respiratory chain protein MT-CO3, encoded by the mitochondrial genome, and the RNA processing factors GRSF1 and LRPPR encoded by the mitochondrial genome. Two-sided permutation t-tests. Each dot is a biological replicate of one MRPL40 null clone analyzed.

**Supplementary Fig. 2.**
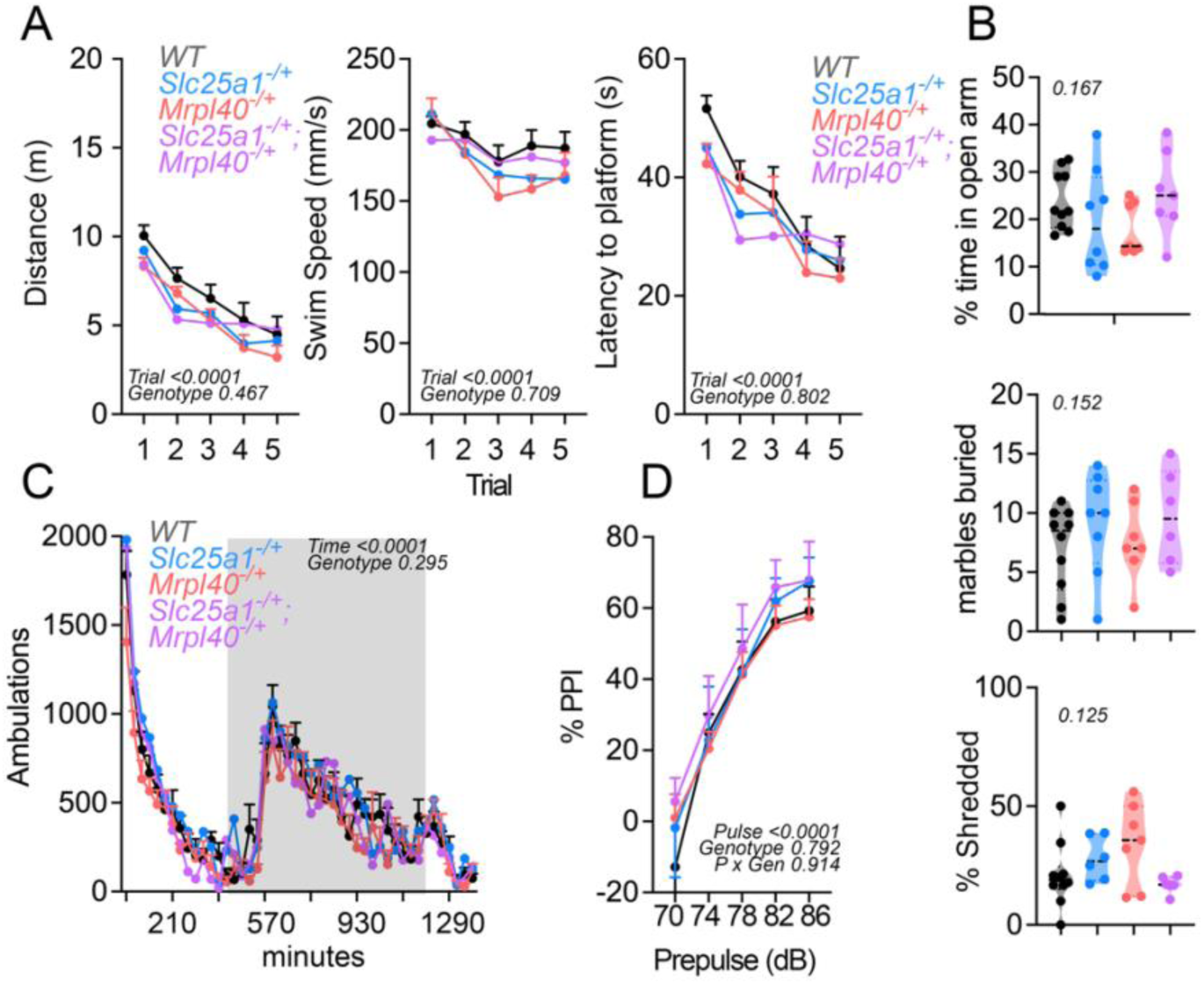
*Mrpl40^-/+^* Female Mice Behavior Analysis. **A**. Swim distance, swim speed, and latency to the platform during acquisition trials of the Morris water maze in 8-week-old females of the indicated genotypes. Two Way Repeated Measures ANOVA n=5-8 per genotype. **B.** Anxiety-like behaviors tests (elevated plus maze, marble burying, and nestlet shredding) are normal in all genotypes tested. Nestlet shredding was analyzed by Kruskal-Wallis test. Other two assays were analyzed by one-way ANOVA. n=6-10 per genotype. **C**. Time course of 23-h locomotor activity of animals of the specified genotype. Grey column denotes dark period. Data represent mean ± SEM. Statistical comparisons were performed using a Two Way Repeated Measures ANOVA with factors genotype, time, and their interactions. n=5-9 per genotype. **D**. Percent PPI, data represent mean ± SEM. Statistical comparisons were performed using a mixed-effects model (Restricted Maximum Likelihood) with factors genotype, prepulse intensity, and their interactions. n=6-10 per genotype.

**Supplementary Fig. 3.**
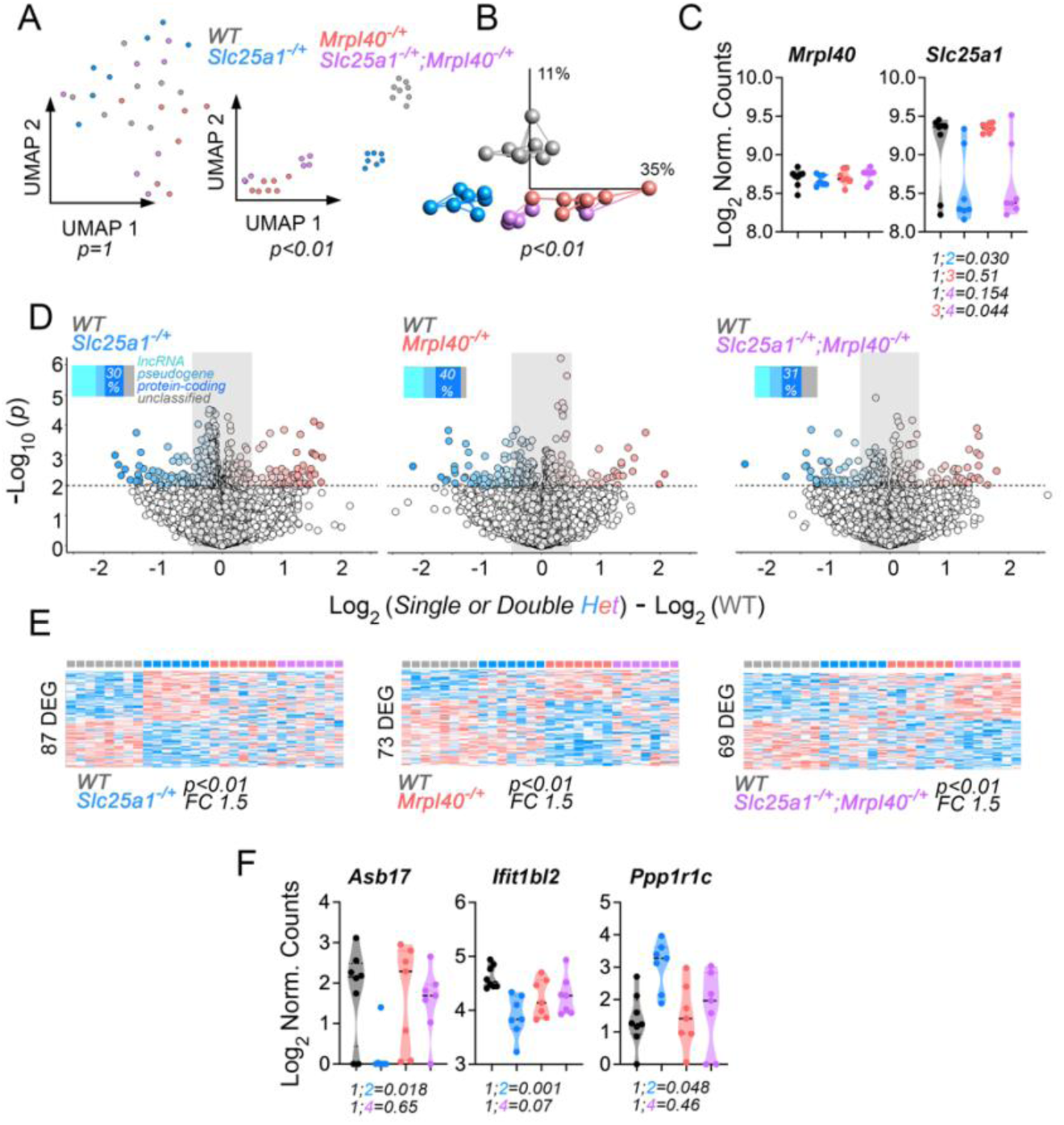
Hippocampal Transcriptome of Single and Double *Slc25a1^-/+^*;*Mrpl40^-/+^* Heterozygotic Mice. **A**. Uniform Manifold Approximation and Projection (UMAP) of hippocampal transcriptome before and after one-way ANOVA thresholding by p<0.01 of the indicated genotype transcriptomes. **B**. PCA of transcriptomes after one-way ANOVA thresholding by p<0.01. Lines represent Euclidean distance clustering. **C.** Normalized hippocampal mRNA counts for Mrpl40 and Slc25a1 transcripts. Kruskal-Wallis test followed by Benjamini and Hochberg multiple corrections. **D**. Volcano plots and **E**. heat maps of paired comparisons between wild type hippocampi from single or transheterozygotic animals. Insets in volcano plots show percentage of diverse types of RNAs differentially expressed. Heat Maps depict z-scored hierarchical clustering of transcripts after thresholding with a cut-off fold of change=1.5 and p<0.01 (Welsh t test). **F** depict violin plots of *Slc25a1^-/+^*-sensitive mRNAs rescued in *Slc25a1^-/+^*;*Mrpl40^-/+^*hippocampus. Kruskal-Wallis test followed by Benjamini and Hochberg multiple corrections. All panels wild type n=8 and all other genotypes n=7.

**Supplementary Table. 1. Transcriptome of Single and Double *Slc25a1^-/+^*;*Mrpl40^-/+^* Heterozygotic Mice after Thresholding (Fold of change=1.5 and p<0.01)**

## Notes

### Competing Interest Statement

The authors have declared no competing interest.

## References

1. Malhotra D, Sebat J. CNVs: harbingers of a rare variant revolution in psychiatric genetics. Cell 2012; 148(6): 1223–1241.

2. Owen MJ, Legge SE, Rees E, Walters JTR, O’Donovan MC. Genomic findings in schizophrenia and their implications. Mol Psychiatry 2023; 28(9): 3638–3647.

3. Pos O, Radvanszky J, Buglyo G, Pos Z, Rusnakova D, Nagy B et al. DNA copy number variation: Main characteristics, evolutionary significance, and pathological aspects. Biomed J 2021; 44(5): 548–559.

4. Willsey HR, Willsey AJ, Wang B, State MW. Genomics, convergent neuroscience and progress in understanding autism spectrum disorder. Nat Rev Neurosci 2022; 23(6): 323–341.

5. Bassett AS, Chow EW. Schizophrenia and 22q11.2 deletion syndrome. Curr Psychiatry Rep 2008; 10(2): 148–157.

6. McDonald-McGinn DM, Sullivan KE, Marino B, Philip N, Swillen A, Vorstman JA, et al. 22q11.2 deletion syndrome. Nat Rev Dis Primers 2015; 1: 15071.

7. Zinkstok JR, Boot E, Bassett AS, Hiroi N, Butcher NJ, Vingerhoets C et al. Neurobiological perspective of 22q11.2 deletion syndrome. Lancet Psychiatry 2019; 6(11): 951–960.

8. Guna A, Butcher NJ, Bassett AS. Comparative mapping of the 22q11.2 deletion region and the potential of simple model organisms. J Neurodev Disord 2015; 7(1): 18.

9. Guo T, Chung JH, Wang T, McDonald-McGinn DM, Kates WR, Hawula W et al. Histone Modifier Genes Alter Conotruncal Heart Phenotypes in 22q11.2 Deletion Syndrome. Am J Hum Genet 2015; 97(6): 869–877.

10. Jensen M, Smolen C, Tyryshkina A, Pizzo L, Sun J, Noss S et al. Genetic modifiers and ascertainment drive variable expressivity of complex disorders. Cell 2025.

11. Smolen C, Girirajan S. The gene dose makes the disease. Cell 2022; 185(16): 2850–2852.

12. Forrest MP, Penzes P. Mechanisms of copy number variants in neuropsychiatric disorders: From genes to therapeutics. Curr Opin Neurobiol 2023; 82: 102750.

13. Iyer J, Singh MD, Jensen M, Patel P, Pizzo L, Huber E et al. Pervasive genetic interactions modulate neurodevelopmental defects of the autism-associated 16p11.2 deletion in Drosophila melanogaster. Nat Commun 2018; 9(1): 2548.

14. Singh MD, Jensen M, Lasser M, Huber E, Yusuff T, Pizzo L et al. NCBP2 modulates neurodevelopmental defects of the 3q29 deletion in Drosophila and Xenopus laevis models. PLoS Genet 2020; 16(2): e1008590.

15. Mlynarski EE, Sheridan MB, Xie M, Guo T, Racedo SE, McDonald-McGinn DM et al. Copy-Number Variation of the Glucose Transporter Gene SLC2A3 and Congenital Heart Defects in the 22q11.2 Deletion Syndrome. Am J Hum Genet 2015; 96(5): 753–764.

16. Bassett AS, Lowther C, Merico D, Costain G, Chow EWC, van Amelsvoort T et al. Rare Genome-Wide Copy Number Variation and Expression of Schizophrenia in 22q11.2 Deletion Syndrome. Am J Psychiatry 2017; 174(11): 1054–1063.

17. Devaraju P, Zakharenko SS. Mitochondria in complex psychiatric disorders: Lessons from mouse models of 22q11.2 deletion syndrome: Hemizygous deletion of several mitochondrial genes in the 22q11.2 genomic region can lead to symptoms associated with neuropsychiatric disease. Bioessays 2017; 39(2).

18. Lujan AL, Foresti O, Wojnacki J, Bigliani G, Brouwers N, Pena MJ et al. TANGO2 is an acyl-CoA binding protein. J Cell Biol 2025; 224(5).

19. Maynard TM, Meechan DW, Dudevoir ML, Gopalakrishna D, Peters AZ, Heindel CC et al. Mitochondrial localization and function of a subset of 22q11 deletion syndrome candidate genes. Mol Cell Neurosci 2008; 39(3): 439–451.

20. Rath S, Sharma R, Gupta R, Ast T, Chan C, Durham TJ et al. MitoCarta3.0: an updated mitochondrial proteome now with sub-organelle localization and pathway annotations. Nucleic Acids Res 2021; 49(D1): D1541–D1547.

21. Wesseling H, Xu B, Want EJ, Holmes E, Guest PC, Karayiorgou M et al. System-based proteomic and metabonomic analysis of the Df(16)A(+/-) mouse identifies potential miR-185 targets and molecular pathway alterations. Mol Psychiatry 2017; 22(3): 384–395.

22. Gokhale A, Hartwig C, Freeman AAH, Bassell JL, Zlatic SA, Sapp Savas C et al. Systems Analysis of the 22q11.2 Microdeletion Syndrome Converges on a Mitochondrial Interactome Necessary for Synapse Function and Behavior. J Neurosci 2019; 39(18): 3561–3581.

23. Harris JJ, Jolivet R, Attwell D. Synaptic energy use and supply. Neuron 2012; 75(5): 762–777.

24. Lopaschuk GD, Ussher JR, Folmes CD, Jaswal JS, Stanley WC. Myocardial fatty acid metabolism in health and disease. Physiol Rev 2010; 90(1): 207–258.

25. Mosaoa R, Kasprzyk-Pawelec A, Fernandez HR, Avantaggiati ML. The Mitochondrial Citrate Carrier SLC25A1/CIC and the Fundamental Role of Citrate in Cancer, Inflammation and Beyond. Biomolecules 2021; 11(2).

26. Ruprecht JJ, Kunji ERS. The SLC25 Mitochondrial Carrier Family: Structure and Mechanism. Trends Biochem Sci 2020; 45(3): 244–258.

27. Brown A, Amunts A, Bai XC, Sugimoto Y, Edwards PC, Murshudov G et al. Structure of the large ribosomal subunit from human mitochondria. Science 2014; 346(6210): 718–722.

28. Greber BJ, Ban N. Structure and Function of the Mitochondrial Ribosome. Annu Rev Biochem 2016; 85: 103–132.

29. Gokhale A, Lee CE, Zlatic SA, Freeman AAH, Shearing N, Hartwig C et al. Mitochondrial Proteostasis Requires Genes Encoded in a Neurodevelopmental Syndrome Locus. J Neurosci 2021; 41(31): 6596–6616.

30. Ohanele C, Peoples JN, Karlstaedt A, Geiger JT, Gayle AD, Ghazal N et al. The mitochondrial citrate carrier SLC25A1 regulates metabolic reprogramming and morphogenesis in the developing heart. Commun Biol 2024; 7(1): 1422.

31. Cheong A, Archambault D, Degani R, Iverson E, Tremblay KD, Mager J. Nuclear-encoded mitochondrial ribosomal proteins are required to initiate gastrulation. Development 2020; 147(10).

32. Tsherniak A, Vazquez F, Montgomery PG, Weir BA, Kryukov G, Cowley GS et al. Defining a Cancer Dependency Map. Cell 2017; 170(3): 564–576 e516.

33. Arafeh R, Shibue T, Dempster JM, Hahn WC, Vazquez F. The present and future of the Cancer Dependency Map. Nat Rev Cancer 2025; 25(1): 59–73.

34. Amici DR, Jackson JM, Truica MI, Smith RS, Abdulkadir SA, Mendillo ML. FIREWORKS: a bottom-up approach to integrative coessentiality network analysis. Life Sci Alliance 2021; 4(2).

35. Bonekamp NA, Larsson NG. SnapShot: Mitochondrial Nucleoid. Cell 2018; 172(1-2): 388–388 e381.

36. Hance N, Ekstrand MI, Trifunovic A. Mitochondrial DNA polymerase gamma is essential for mammalian embryogenesis. Hum Mol Genet 2005; 14(13): 1775–1783.

37. Larsson NG, Wang J, Wilhelmsson H, Oldfors A, Rustin P, Lewandoski M et al. Mitochondrial transcription factor A is necessary for mtDNA maintenance and embryogenesis in mice. Nat Genet 1998; 18(3): 231–236.

38. Oetting WS. The tyrosinase gene and oculocutaneous albinism type 1 (OCA1): A model for understanding the molecular biology of melanin formation. Pigment Cell Res 2000; 13(5): 320–325.

39. Thermann R, Neu-Yilik G, Deters A, Frede U, Wehr K, Hagemeier C et al. Binary specification of nonsense codons by splicing and cytoplasmic translation. EMBO J 1998; 17(12): 3484–3494.

40. Choquet C, Kelly RG, Miquerol L. Defects in Trabecular Development Contribute to Left Ventricular Noncompaction. Pediatr Cardiol 2019; 40(7): 1331–1338.

41. Liu Y, Chen H, Shou W. Potential Common Pathogenic Pathways for the Left Ventricular Noncompaction Cardiomyopathy (LVNC). Pediatr Cardiol 2018; 39(6): 1099–1106.

42. Imes S, Parker DA, Chen M, Cubells JF, Walker EF, Duncan E. The acoustic startle response in 22q11 deletion syndrome: from animal models to humans. Front Neurosci 2025; 19: 1630109.

43. Lindsay EA, Botta A, Jurecic V, Carattini-Rivera S, Cheah YC, Rosenblatt HM et al. Congenital heart disease in mice deficient for the DiGeorge syndrome region. Nature 1999; 401(6751): 379–383.

44. Paylor R, Glaser B, Mupo A, Ataliotis P, Spencer C, Sobotka A et al. Tbx1 haploinsufficiency is linked to behavioral disorders in mice and humans: implications for 22q11 deletion syndrome. Proc Natl Acad Sci U S A 2006; 103(20): 7729–7734.

45. Merscher S, Funke B, Epstein JA, Heyer J, Puech A, Lu MM et al. TBX1 is responsible for cardiovascular defects in velo-cardio-facial/DiGeorge syndrome. Cell 2001; 104(4): 619–629.

46. Yagi H, Furutani Y, Hamada H, Sasaki T, Asakawa S, Minoshima S et al. Role of TBX1 in human del22q11.2 syndrome. Lancet 2003; 362(9393): 1366–1373.

47. Eom TY, Schmitt JE, Li Y, Davenport CM, Steinberg J, Bonnan A et al. Tbx1 haploinsufficiency leads to local skull deformity, paraflocculus and flocculus dysplasia, and motor-learning deficit in 22q11.2 deletion syndrome. Nat Commun 2024; 15(1): 10510.

48. Jensen M, Girirajan S. An interaction-based model for neuropsychiatric features of copy-number variants. PLoS Genet 2019; 15(1): e1007879.

49. Meisel JD, Miranda M, Skinner OS, Wiesenthal PP, Wellner SM, Jourdain AA et al. Hypoxia and intra-complex genetic suppressors rescue complex I mutants by a shared mechanism. Cell 2024; 187(3): 659–675 e618.

50. Wynne ME, Lane AR, Singleton KS, Zlatic SA, Gokhale A, Werner E et al. Heterogeneous Expression of Nuclear Encoded Mitochondrial Genes Distinguishes Inhibitory and Excitatory Neurons. eNeuro 2021; 8(4).

51. Devaraju P, Yu J, Eddins D, Mellado-Lagarde MM, Earls LR, Westmoreland JJ et al. Haploinsufficiency of the 22q11.2 microdeletion gene Mrpl40 disrupts short-term synaptic plasticity and working memory through dysregulation of mitochondrial calcium. Mol Psychiatry 2017; 22(9): 1313–1326.

52. Fernandez A, Meechan DW, Karpinski BA, Paronett EM, Bryan CA, Rutz HL et al. Mitochondrial Dysfunction Leads to Cortical Under-Connectivity and Cognitive Impairment. Neuron 2019; 102(6): 1127–1142 e1123.

53. Campbell PD, Lee I, Thyme S, Granato M. Mitochondrial proteins encoded by the 22q11.2 neurodevelopmental locus regulate neural stem and progenitor cell proliferation. Mol Psychiatry 2023; 28(9): 3769–3781.

54. Sprecher E, Ishida-Yamamoto A, Mizrahi-Koren M, Rapaport D, Goldsher D, Indelman M et al. A mutation in SNAP29, coding for a SNARE protein involved in intracellular trafficking, causes a novel neurocutaneous syndrome characterized by cerebral dysgenesis, neuropathy, ichthyosis, and palmoplantar keratoderma. Am J Hum Genet 2005; 77(2): 242–251.

55. Delprato A, Xiao E, Manoj D. Connecting DCX, COMT and FMR1 in social behavior and cognitive impairment. Behav Brain Funct 2022; 18(1): 7.

56. Gogos JA, Santha M, Takacs Z, Beck KD, Luine V, Lucas LR et al. The gene encoding proline dehydrogenase modulates sensorimotor gating in mice. Nat Genet 1999; 21(4): 434–439.

57. Papaleo F, Crawley JN, Song J, Lipska BK, Pickel J, Weinberger DR et al. Genetic dissection of the role of catechol-O-methyltransferase in cognition and stress reactivity in mice. J Neurosci 2008; 28(35): 8709–8723.

58. Stark KL, Burt RA, Gogos JA, Karayiorgou M. Analysis of prepulse inhibition in mouse lines overexpressing 22q11.2 orthologues. Int J Neuropsychopharmacol 2009; 12(7): 983–989.

59. LaMantia AS. Polygenicity in a box: Copy number variants, neural circuit development, and neurodevelopmental disorders. Curr Opin Neurobiol 2024; 89: 102917.

60. Ishikawa K, Miyata D, Hattori S, Tani H, Kuriyama T, Wei FY et al. Accumulation of mitochondrial DNA with a point mutation in tRNA(Leu(UUR)) gene induces brain dysfunction in mice. Pharmacol Res 2024; 208: 107374.

61. Pietropaolo S, Crusio WE. Strain-dependent changes in acoustic startle response and its plasticity across adolescence in mice. Behav Genet 2009; 39(6): 623–631.

62. Shoji H, Takao K, Hattori S, Miyakawa T. Age-related changes in behavior in C57BL/6J mice from young adulthood to middle age. Mol Brain 2016; 9: 11.

63. Paylor R, Crawley JN. Inbred strain differences in prepulse inhibition of the mouse startle response. Psychopharmacology (Berl) 1997; 132(2): 169–180.

64. Paylor R, McIlwain KL, McAninch R, Nellis A, Yuva-Paylor LA, Baldini A et al. Mice deleted for the DiGeorge/velocardiofacial syndrome region show abnormal sensorimotor gating and learning and memory impairments. Hum Mol Genet 2001; 10(23): 2645–2650.

65. Sumitomo A, Horike K, Hirai K, Butcher N, Boot E, Sakurai T et al. A mouse model of 22q11.2 deletions: Molecular and behavioral signatures of Parkinson’s disease and schizophrenia. Sci Adv 2018; 4(8): eaar6637.

66. Scarborough J, Mueller F, Weber-Stadlbauer U, Richetto J, Meyer U. Dependency of prepulse inhibition deficits on baseline startle reactivity in a mouse model of the human 22q11.2 microdeletion syndrome. Genes Brain Behav 2019; 18(4): e12523.

67. Didriksen M, Fejgin K, Nilsson SR, Birknow MR, Grayton HM, Larsen PH, et al. Persistent gating deficit and increased sensitivity to NMDA receptor antagonism after puberty in a new mouse model of the human 22q11.2 microdeletion syndrome: a study in male mice. J Psychiatry Neurosci 2017; 42(1): 48–58.

68. Saito R, Koebis M, Nagai T, Shimizu K, Liao J, Wulaer B, et al. Comprehensive analysis of a novel mouse model of the 22q11.2 deletion syndrome: a model with the most common 3.0-Mb deletion at the human 22q11.2 locus. Transl Psychiatry 2020; 10(1): 35.

69. Saito R, Miyoshi C, Koebis M, Kushima I, Nakao K, Mori D et al. Two novel mouse models mimicking minor deletions in 22q11.2 deletion syndrome revealed the contribution of each deleted region to psychiatric disorders. Mol Brain 2021; 14(1): 68.

70. Hiramoto T, Sumiyoshi A, Kato R, Yamauchi T, Takano T, Kang G et al. Highly demarcated structural alterations in the brain and impaired social incentive learning in Tbx1 heterozygous mice. Mol Psychiatry 2025; 30(5): 1876–1886.

71. Caterino M, Paris D, Torromino G, Costanzo M, Flore G, Tramice A et al. Brain and behavioural anomalies caused by Tbx1 haploinsufficiency are corrected by vitamin B12. Life Sci Alliance 2025; 8(2).

72. Swerdlow NR, Geyer MA, Braff DL. Neural circuit regulation of prepulse inhibition of startle in the rat: current knowledge and future challenges. Psychopharmacology (Berl) 2001; 156(2-3): 194–215.

73. Rooney JP, Ryde IT, Sanders LH, Howlett EH, Colton MD, Germ KE et al. PCR based determination of mitochondrial DNA copy number in multiple species. Methods Mol Biol 2015; 1241: 23–38.

74. Bankhead P, Loughrey MB, Fernandez JA, Dombrowski Y, McArt DG, Dunne PD et al. QuPath: Open source software for digital pathology image analysis. Sci Rep 2017; 7(1): 16878.

75. Andrews S. FastQC A Quality Control tool for High Throughput Sequence Data. Version 0.12.0 edn2023.

76. Afgan E, Baker D, Batut B, van den Beek M, Bouvier D, Cech M et al. The Galaxy platform for accessible, reproducible and collaborative biomedical analyses: 2018 update. Nucleic Acids Res 2018; 46(W1): W537–W544.

77. Kim D, Langmead B, Salzberg SL. HISAT: a fast spliced aligner with low memory requirements. Nat Methods 2015; 12(4): 357–360.

78. Liao Y, Smyth GK, Shi W. featureCounts: an efficient general purpose program for assigning sequence reads to genomic features. Bioinformatics 2014; 30(7): 923–930.

79. Love MI, Huber W, Anders S. Moderated estimation of fold change and dispersion for RNA-seq data with DESeq2. Genome Biol 2014; 15(12): 550.

80. Ho J, Tumkaya T, Aryal S, Choi H, Claridge-Chang A. Moving beyond P values: data analysis with estimation graphics. Nat Methods 2019; 16(7): 565–566.

81. Kuleshov MV, Jones MR, Rouillard AD, Fernandez NF, Duan Q, Wang Z et al. Enrichr: a comprehensive gene set enrichment analysis web server 2016 update. Nucleic Acids Res 2016; 44(W1): W90–97.

82. Antonicka H, Lin ZY, Janer A, Aaltonen MJ, Weraarpachai W, Gingras AC et al. A High-Density Human Mitochondrial Proximity Interaction Network. Cell Metab 2020; 32(3): 479–497 e479.

